# Eukaryotic genomes from a global metagenomic dataset illuminate trophic modes and biogeography of ocean plankton

**DOI:** 10.1101/2021.07.25.453713

**Authors:** Harriet Alexander, Sarah K. Hu, Arianna I. Krinos, Maria Pachiadaki, Benjamin J. Tully, Christopher J. Neely, Taylor Reiter

## Abstract

Metagenomics is a powerful method for interpreting the ecological roles and physiological capabilities of mixed microbial communities. Yet, many tools for processing metagenomic data are not designed to consider eukaryotes, nor are they built for an increasing amount of sequence data. EukHeist is an automated pipeline to retrieve eukaryotic and prokaryotic metagenome assembled genomes (MAGs) from large-scale metagenomic datasets. We developed the EukHeist workflow to specifically process large amounts of both metagenomic and/or metatranscriptomic sequence data in an automated and reproducible fashion. Here, we applied EukHeist to the large-size fraction data (0.8-2000*µm*) from *Tara* Oceans to recover both eukaryotic and prokaryotic MAGs, which we refer to as TOPAZ (*Tara* Oceans Particle-Associated MAGs). The TOPAZ MAGs consisted of >900 environmentally-relevant eukaryotic MAGs and >4,000 bacterial and archaeal MAGs. The bacterial and archaeal TOPAZ MAGs expand the known marine phylogenetic diversity through the increase in coverage of likely particle- and host-associated taxa. We also demonstrate an approach to infer the putative functional mode of the recovered eukaryotic MAGs. A global survey of the TOPAZ MAGs enabled the identification of ecological cohorts, driven by specific environmental factors, and putative host-microbe associations.

**Importance:** Despite the ecological importance of single-celled eukaryotic organisms in marine environments, the majority are difficult to cultivate in the lab. Sequencing genetic material extracted from environmental samples enables researchers to document naturally-occurring protistan communities. However, conventional sequencing methodologies cannot separate out the genomes of individual organisms. To more completely capture the entire genomic content of mixed protistan community, we can create bins of sequences that represent the same organism. We developed a pipeline that enables scientists to bin individual organisms out of metagenomic reads, and show results that provide exciting insights into what protistan communities are present in the ocean and what roles they play in the ecosystem. Here, a global survey of both eukaryotic and prokaryotic MAGs enabled the identification of ecological cohorts, driven by specific environmental factors, and putative host-microbe associations. Accessible and scalable computational tools, such as EukHeist, are likely to accelerate the identification of meaningful genetic signatures from large datasets, ultimately expanding the eukaryotic tree of life.

## Introduction

Unicellular microbial eukaryotes, or protists, play a critical part in many ecosystems found on the planet. In addition to their vast morphological and taxonomic diversity, protists exhibit a range of functional roles and trophic strategies (*1*). Protists are centrally important to global biogeochemical cycles, mediating the pathways for the synthesis and processing of carbon and nutrients in the environment (*2–4*). Despite their importance across ecosystems and in the global carbon cycle, research on microbial eukaryotes typically lags behind that of bacteria and archaea (*5, 6*). Consequently, fundamental questions surrounding microbial eukaryotic ecological function *in situ* remain unresolved. Novel approaches that enable genome retrieval from meta’omic data provide a means of bridging that knowledge gap.

Assembled genetic fragments (derived from metagenomic reads) can be grouped together based on their abundances, co-occurrences, and tetranucleotide frequency to reconstruct likely genomic collections, often called bins (*7–10*). These bins can be further refined through a series of steps to ultimately represent metagenome assembled genomes or MAGs (*11–14*). Binning metagenomic data into MAGs has revolutionized how researchers ask questions about microbial communities and has enabled the identification of novel bacterial and archaeal taxa and functional traits (*15, 16*), but only recently have the recovery of eukaryotic MAGs become more common (*17–19*). The reason for this is arguably twofold: (1) eukaryotic genomic complexity (*20*) complicates both metagenome assembly and MAG retrieval; and (2) there is a bias in currently available metagenomic computational tools towards the study of bacterial and archaeal members of the community. Much can be learned about the diversity and role of eukaryotes in our environment from eukaryotic MAG retrieval (*21*).

Here we developed and applied EukHeist, a scalable and reproducible pipeline to facilitate the reconstruction, taxonomic assignment, and annotation of prokaryotic and eukaryotic metagenome assembled genomes (MAGs) from mixed community metagenomes. The EukHeist pipeline was applied to a metagenomic dataset from the *Tara* Oceans expedition protist-size fractions samples (*22*), which encompasses more than 20Tb of raw sequence data. From large-size fraction metagenomic samples, over 4,000 prokaryotic MAGs and 900 eukaryotic MAGs were recovered, representing a multi-domain approach for MAG retrieval for mixed microbial communities.

## Results and Discussion

The EukHeist metagenomic pipeline was designed to automate the recovery and classification of eukaryotic and prokaryotic MAGs from large-scale environmental metagenomic datasets. EukHeist was applied to the metagenomic data from the large-size fraction metagenomic samples (0.8-2000*µm*) from *Tara* Oceans (*22*), which is dominated by eukaryotic organisms. We generated 94 co-assembled metagenomes based on the ocean region, size fraction, and depth of the samples (Figure S1), which totaled 180 Gbp in length (Supplementary Table 1). A total of 988 eukaryotic MAGs and 4,022 prokaryotic MAGs were recovered; these MAGs have been made available under the name *Tara* Oceans Particle Associated MAGs, or TOPAZ (Supplementary Tables 2 and 3). The TOPAZ MAGs expand the current repertoire of publicly available eukaryotic genomic references for the marine environment and complement other efforts to recover eukaryotic MAGs from the same large size-fraction data set (*17*). Here, we use the information gained from a highly reproducible and automated approach to explore questions related to the biogeographical distribution and functional potential of eukaryotic marine communities.

### Eukaryotic genome recovery from metagenomes covers major eukaryotic supergroups

The EukHeist classification pipeline identified 988 putative eukaryotic MAGs following the refinement of recovered metagenomic bins based on length (*>* 2.5 Mbp) and proportion of base pairs predicted to be eukaryotic in origin by EukRep (*23*) (Figure S4). Protein coding regions in the eukaryotic MAGs were predicted using the EukMetaSanity pipeline (*24*), and the likely taxonomic assignment of each bin was made with MMSeqs (*25*) and EUKulele (*26*) (Supplementary Table 2). Of the 988 eukaryotic MAGs recovered, 713 MAGs were estimated to be more than 10% complete based on the presence of core eukaryotic BUSCO orthologs (*27*). For the purposes of our subsequent analyses, we only consider the highly complete eukaryotic TOPAZ MAGs, or those that were greater than 30% complete based on BUSCO ortholog presence (n=485) (Figure 1).

**Figure 1.**
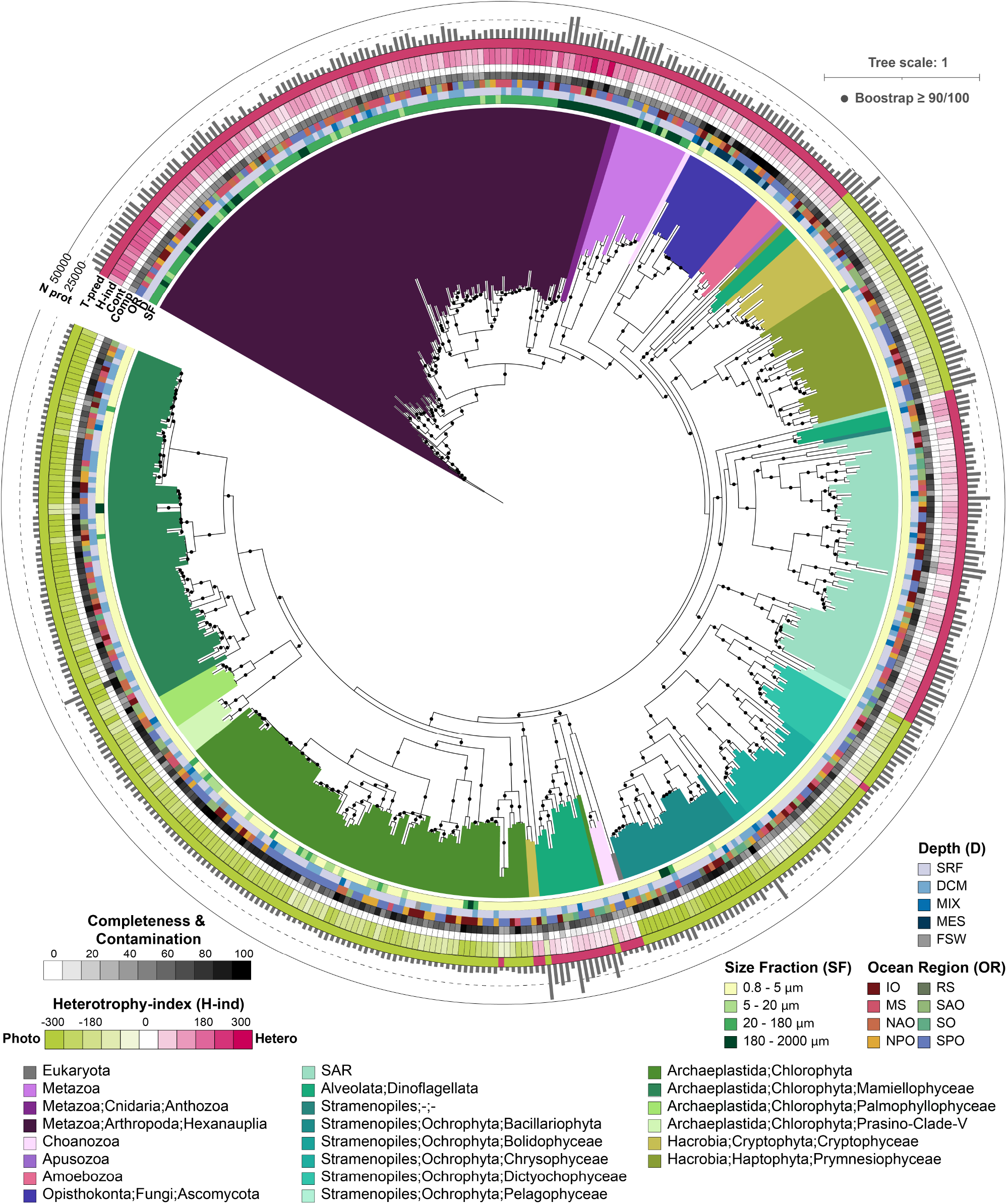
TOPAZ eukaryotic MAGs span the eukaryotic tree of life. The maximum likelihood tree was inferred from a concatenated protein alignment of 49 proteins from the eukaryotic BUSCO gene set that were found to be commonly present across at least 75% of the 485 TOPAZ eukaryotic MAGs that were estimated to be >30% complete based on BUSCO ortholog presence. The MAG names were omitted but the interactive version of the tree containing the MAG names can be accessed through iTOL (https://itol.embl.de/shared/halexand). Branches (nodes) are colored based on consensus protein annotation estimated by EUKulele and MMSeqs. The Ocean Region (OR), Depth (D), and Size Fraction (SF) of the co-assembly that a MAG was isolated from is color coded as colored bars. The completeness (comp) and contamination (cont) as estimated based on BUSCO presence are depicted as a heatmap. Predicted Heterotrophy Index (H-index), which ranges from phototroph-like (−300) to heterotroph-like (300) is shown as a heatmap. The predicted trophic mode (T-pred) based on the trophy random forest classifier with heterotroph (pink) and phototroph (green), is depicted. The number of proteins predicted with EukMetaSanity are shown as a bar graph along the outermost ring.

Eukaryotic genomes are known to be both larger and have higher proportions of non-coding DNA than bacterial genomes (*20*). On average across sequenced eukaryotic genomes, 33.1% of genomic content codes for genes (2.6% - 59.8% for the 1st and 3rd quartiles), while bacterial genomes have a higher proportion of coding regions (86.9%; 83.9% - 89.3%) (*28*). The high-completion TOPAZ eukaryotic MAGs have an average of 73.7% ± 14.3% gene coding regions (Figure S9). This trend of a higher proportion of coding regions was consistent across eukaryotic groups, where Haptophyta and Ochrophyta TOPAZ MAGs had an average coding region of 80.3 ± 4.9% and 78.1 ± 6.3%, respectively. Genomes from cultured Haptophyta (*Emiliania huxleyi* CCMP1516 with 31 Mb or 21.9% (*29*)), and Ochrophyta (*Phaeodactylum tricornutum* with 15.4 Mb or 57.3% (*30*)) had significantly lower proportions of protein coding regions within their genomes compared to TOPAZ MAGs. The lowest percentages of gene coding were within Metazoan and Fungal TOPAZ MAGs, with 52.6 ± 9.8% and 58.8 ± 6.7%, respectively. As a point of comparison, the human genome is estimated to have ≈ 34 Mb or ≈ 1.2% of the genome coding for proteins (*31*). Globally, the higher gene coding percentages for the recovered eukaryotic TOPAZ MAGs likely reflect biases caused by the use of tetranucleotide frequencies in the initial binning (*9*) as well as challenges inherent in the assembly of non-coding and repeat-rich regions of eukaryotic genomes.

In order to evaluate the taxonomic breadth represented in the TOPAZ MAGs, estimated taxonomy of each MAG based on protein-consensus annotation was used for phylogenetic placement of TOPAZ MAGs (Figure 1). The recovered MAGs spanned 8 major eukaryotic supergroups: Archaeplastida (Chlorophyta), Opisthokonta (Metazoa, Choanoflagellata, and Fungi), Amoebozoa, Apusozoa, Haptista (Haptophyta), Cryptista (Cryptophyta), and the SAR supergroup (Stramenopiles, Alveolata, and Rhizaria) (*32*), similar to other eukaryotic MAG recovery efforts from the Tara Oceans dataset (Figure S14) (*17*). Eukaryotic MAGs were retrieved from all ocean regions surveyed, with the largest number of high-completion TOPAZ MAGs recovered from the South Pacific Ocean Region (SPO) (n=143) and the fewest recovered from the Southern Ocean (SO) (n=11) and Red Sea (RS) (n=12) (Figure S8). These regional trends in MAG recovery and taxonomy aligned with the overall sequencing depth at each of these locations (Supplementary Table 1), with fewer, less diverse MAGs recovered from the SO and RS (Figures S5 and S8).

The largest number of MAGs was recovered from the smallest size fraction (0.8 − 5*µm*) (n=311) (Figures 1 and S5), and yielded the highest taxonomic diversity, including MAGs from all the major supergroups listed above (Figure S5). The groups which made up the largest proportion of small size fraction MAGs were Chlorophyta (n=133), Ochrophyta (n=33), or taxa likely to be within the SAR group (Stramenopiles, Alveolata, and Rhizaria; n=56). Chlorophyta MAGs were smaller and had fewer predicted proteins relative to other eukaryotic MAGs, despite demonstrating comparable completeness metrics; the average Chlorophyta MAG size was 13.9 Mbp with 7525 predicted proteins (Figures S6 and S9). By contrast, Cryptophyta and Haptophyta had the largest average MAG size with 50.8 Mbp and 44.4 Mbp with an average of 23500 and 24400 predicted proteins, respectively (Figure S9). Fewer eukaryotic MAGs were recovered from the other size fractions 5 − 20*µm* (n=20), 20 − 180*µm* (n=87), and 180−2000*µm* (n=39) (Figure S5), instead these larger size fractions recovered a higher total number of metazoan MAGs. Metazoan MAGs had the lowest average completeness (50 ± 13%) (Figures S7 and S9); where the average size of recovered metazoan MAGs was 43.2 Mb (6.5-177Mbp), encompassing an average of 14600 proteins (Figure S9). 76 of the 123 metazoan MAGs likely belong to the Hexanauplia (Copepoda) class; copepod genomes have been estimated to be up to 2.5 Gb with high variation (10-fold difference) across sequenced members (*33*).

MAGs were also retrieved from all discrete sampling depths: surface, SRF (n=315), deep chlorophyll max, DCM (n=133), mesopelagic, MES (n=13), as well as samples with no discrete depth, MIX (n=21) and the filtered seawater controls, FSW (n=3). Notably, the FSW included 1 Chlorophyta MAG (TOPAZ_IOF1_E003) that was estimated to be 100% complete with no contamination (Supplementary Table 2). These results suggest that variables such as *in situ* diversity, cell size (and genomic size), and sampling protocols influence our ability to obtain high quality and highly complete eukaryotic MAGs.

The composition of TOPAZ MAGs from basin-scale mesopelagic co-assemblies recovered a higher percentage of fungi relative to other depths. This is similar to other mesopelagic and bathypelagic molecular surveys, where the biomass of fungi is thought to outweigh other eukaryotes (*34–36*). Further, fungal MAGs had the highest overall average completeness (87 ± 15%) (Figures S7 and S9).

A total of 16 highly complete fungal MAGs were also recovered, of those, 11 originated from the MES (Figures 1 and S8). Putative fungal TOPAZ MAGs were recovered from the phyla Ascomycota (n=10) and Basidiomycota (n=1) and ranged in size from 12.5-47.8 Mb (Figure S9), which are within range of known average genome sizes for these groups, 36.9 and 46.5 Mb, respectively (*37*).

The metagenomic read recruitment to these TOPAZ MAGs paralleled MAG recovery, where metazoan MAGs dominated the larger size fractions (20 − 180*µm* and 180 − 2000*µm*) across both the surface and DCM for all stations, and Chlorophyta MAGs were dominant across most of the small size fraction stations (0.8 − 5*µm*) (Figure S11). A notable exception were the stations from the Southern Ocean, where Haptophyta and Ochrophyta were most abundant in all size fractions. Compared to the samples from the photic zone, SRF and DCM, the average recruitment of reads from the MES was far lower (24500 ± 34450 average CPM in the MES compared to 131000 ± 104000 and 136000 ± 85000 for the SRF and DCM, respectively Figure S11). This suggests that MAG binning did not capture the complete eukaryotic community, which can be partly explained by low read recruitment from the highly diverse and distinct microbial populations across different mesopelagic samples (*35*). Alternatively, this might suggest that the communities sampled were dominated by prokaryotic biomass (*38*).

### Prediction of trophic mode from MAG gene content

Eukaryotic microbes can exhibit a diversity of functional traits and trophic strategies in the marine environment (*1, 39*), including phototrophy, heterotrophy, and mixotrophy. Phototrophic protists are responsible for a significant fraction of the organic carbon synthesis via primary production; these phototrophs dominate the microbial biomass and diversity in the sunlit layer of the oceans (*39, 40*). Phagotrophic protists (heterotrophs), which ingest bacteria, archaea, and smaller eukaryotes, and parasitic protists are known to account for a large percentage of mortality in food webs (*1,39,41*). Protists are also capable of mixed nutrition (mixotrophy), where a single-cell exhibits a combination of phototrophy and heterotrophy (*42*). Typically, the identification of trophic mode has relied upon direct observations of isolates within an lab setting, with more recent efforts including transcriptional profiling as a means of assessing trophic strategy (*43, 44*). Scaling up these culture-based observations to environmentally-relevant settings (*45–47*) has been an important advance in the field for exploring complex communities without cultivation. An outcome from these studies has been the realization that trophic strategies are not governed by single genes (*48*); in reality, trophic strategy will be shaped by an organisms’ physiological potential and environmental setting. Therefore larger genomic and transcriptomic efforts to predict or characterize presumed trophic strategies among mixed microbial communities will greatly contribute to our understanding of the role that microorganisms play in global biogeochemical cycles, by enabling the observation of functional traits and strategies *in situ*.

Large scale meta’omic results, such the TOPAZ MAGs recovered here, can be leveraged alongside presently available reference data to enable the prediction of biological traits (such as trophic mode) without *a priori* information. Machine learning (ML) applications can be implemented to access the potential of these large datasets. ML approaches have been recently shown to be capable of accurate functional prediction and cell type annotation using genetic input, in particular for cancer cell prediction (*49–51*), and functional gene and phenotype prediction in plants (*52*). Recently, these approaches have been applied to culture and environmental transcriptomic data to predict trophic mode using currently available trophy annotations (*53–55*). Here, we apply an independent machine learning model to the eukaryotic TOPAZ MAGs to predict each organisms’ capacity for various metabolisms.

Using a reference set built from protistan transcriptomic data, we predicted the trophic mode of the highly complete TOPAZ MAGs using machine learning and direct estimation via presence of important KEGG pathways (eq. (8)). As the gradient of trophic mode among protists is not strictly categorical, we calculated a Heterotrophy Index (H-index) that places the TOPAZ MAGs on a scale of highly phototrophic (negative values) to highly heterotrophic (positive values) (Figures 1 and 2). Thus, for all sufficiently complete (≥ 30%) TOPAZ MAGs we have predicted both a gross trophic category (heterotrophic (*n* = 227), mixotrophic (*n* = 0), or phototrophic (*n* = 258) as well as the quantitative extent of heterotrophy (H-index, eq. (8)). Broadly, the trophic predictions aligned well with the putative taxonomy of each MAG (Figures 1 and 2). For example, TOPAZ MAGs that had taxonomic annotation of well known heterotrophic lineages (Metazoa, Fungi), were predicted as heterotrophs based on our model. Further, our data-driven trophic mode predictions correlate well with an independent model designed to identify the presence of photosynthetic machinery and capacity for phagotrophy (*53, 54*) (Figures S19 and S20).

**Figure 2.**
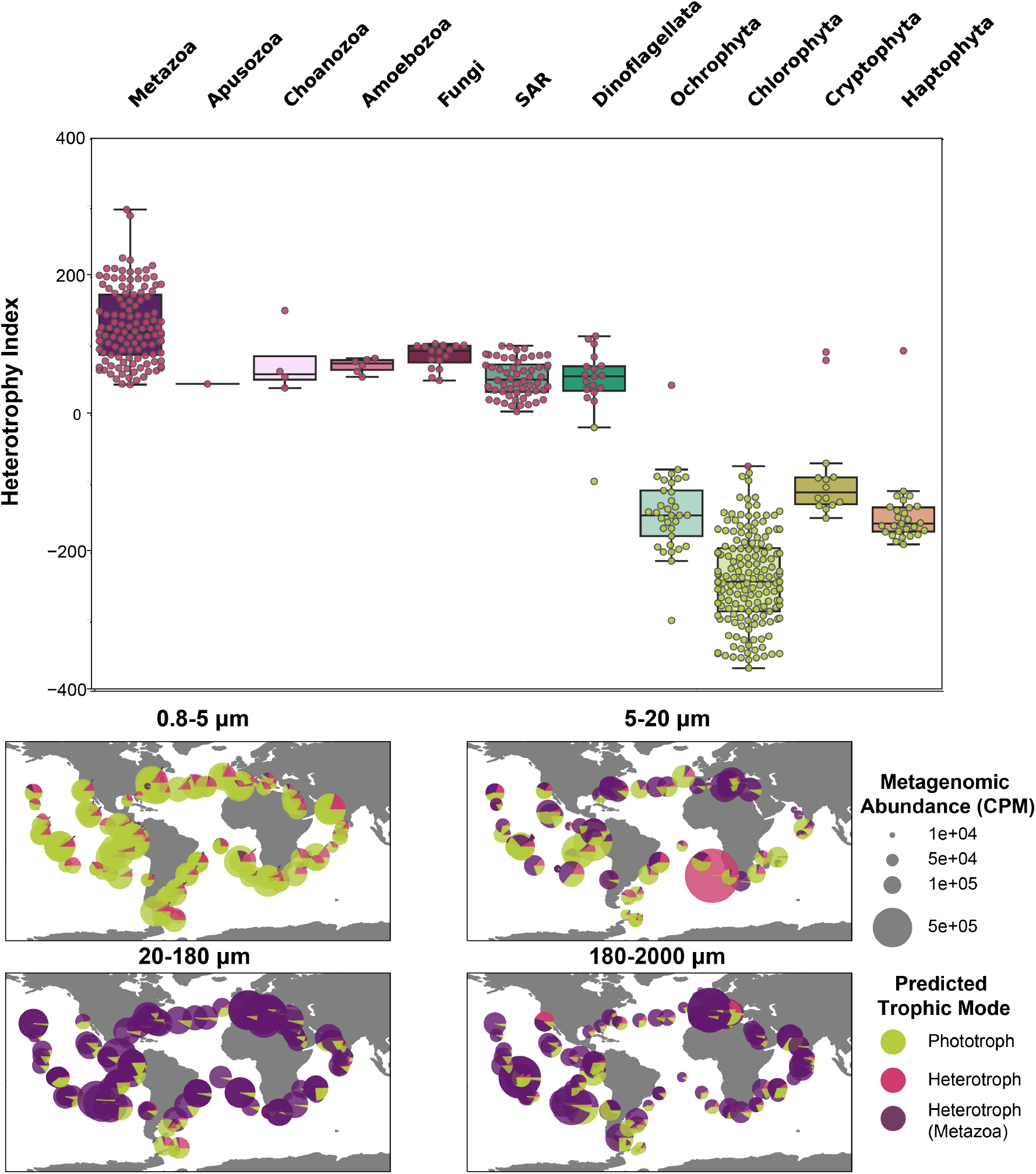
Estimated trophic status of TOPAZ eukaryotic MAGs. (Top) Trophic status was predicted for each high-completion TOPAZ eukaryotic MAG using a Random Forest model trained on the presence and absence of KEGG orthologs and is shown as a color (green, phototroph, pink, heterotroph). The Heterotrophy Index (H-index) (Equation (8)) for each MAG is plotted with a box plot showing the range of the H-index for each higher level group. (Bottom) The relative distribution and abundance of Phototroph (green), non-Metazoan Heterotroph (Pink), and Metazoan Heterotroph (Purple) is depicted across all surface samples. Plots are subdivided by size classes. ‘SAR’ denotes MAGs with taxonomy assignments that were not resolved beyond the SAR supergroup (Stramenopile, Alveolate, or Rhizaria).

Despite evidence that many lineages recovered include known mixotrophs, no TOPAZ MAGs were identified as mixotrophic using this approach. Instead, the utility of the H-index enables us to still consider mixotrophic-capable MAGs. We explore the likely reasons for this more deeply in the Section 2.3, but one potential explanation is that MAG recovery targets the genomic content of a eukaryotic lineage and the evolutionary history of phototrophy and heterotrophy is complicated and varies with respect to species (*56*). Therefore, the genetic composition of MAGs may reflect encoded metabolisms that are not necessarily exhibited *in situ*. Mixotrophy is not a singular trait, but rather a spectrum of metabolic abilities that are largely driven by the microorganisms’ nutritional needs and surrounding environment.

This work demonstrates the value of large untargeted genetic approaches to gain insight into the *in situ* metabolisms of less explored branches of the eukaryotic tree of life. Automated recovery of eukaryotic MAGs, independent of a reference database, and the trophic mode prediction demonstrates how we can begin to parse the metabolic contributions of individual eukaryotes to mixed microbial communities. While we cannot confidently annotate beyond specific taxonomic levels or protein identities, our ML model approaches still allows us to capture predicted nutritional strategies alongside the environmental context provided by the large-scale global sampling effort. Continued culturing efforts combined with large-scale meta’omic studies will continue to improve such ML models focused on complex traits and ultimately our ability to predict trophic mode. We suggest that the integration of metagenomic and metatranscriptomic datasets might better reflect the active strategies being used.

### TOPAZ prokaryotic MAGs distinct from previous marine MAG recovery efforts

The vast majority of the retrieved prokaryotic MAGs belonged to Bacteria. High-quality non-redundant TOPAZ (HQ-NR-TOPAZ) MAGs were comprised of 711 bacterial and 5 archaeal MAGs belonging to 30 different phyla (Figure 3 and Supplementary Table 4); an additional 15 phyla were recovered in the medium quality (MQ) MAGs. Of the 716 HQ-NR-TOPAZ MAGs, 507 were unique based on a 99% ANI comparison threshold with MAGs generated from previous binning efforts from *Tara* Oceans metagenomic data, including Delmont et al. (*12*) (TARA), Tully et al. (*13*) (TOBG), and Parks et al. (*11*) (UBA) (Figure 3). The phylogenetic diversity captured by the TOPAZ MAGs was quantified by a comparison to a “neutral” reference set of genomes; these neutral references approximate the state of marine microbial genomes, dominated by isolate genomes, previous to the incorporation of the *Tara* Oceans-derived MAGs (Table 1). Relative to the neutral genomic references, the entire TOPAZ NR (includes both HQ and MQ) set represented a 42.8% phylogenetic gain (as measured by additional branch length contributed by a set of data) and 59.9% phylogenetic diversity (as measured by the total branch length spanned by a set of taxa), as compared to efforts focused solely on the smaller size fractions such as TARA and UBA, which had a smaller degree of gain (31.0% and 25.8%, respectively) and diversity (44.4% and 40.5%, respectively) (Table 1). An inclusive tree containing the neutral reference and all *Tara* Oceans MAGs (TOBG + UBA + TARA + TOPAZ), the TOPAZ NR MAGs represented 14.4% of the phylogenetic gain and 44.7% phylogenetic diversity, suggesting that the TOPAZ MAGs offer the largest increase in phylogenetic novelty when compared to MAGs reconstructed from the metagenomes of the smaller size fractions (*<* 5.00*µm*). The TOPAZ MAGs primarily originated from the larger *Tara* Ocean size fraction samples, and thus include a higher proportion of more complex host-and particle-associated bacterial communities. The novelty of the HQ-and MQ-NR-TOPAZ MAGs here, suggests that these particle-associated MAGs are overlooked and current genome databases are largely skewed towards free-living bacteria.

**Figure 3.**
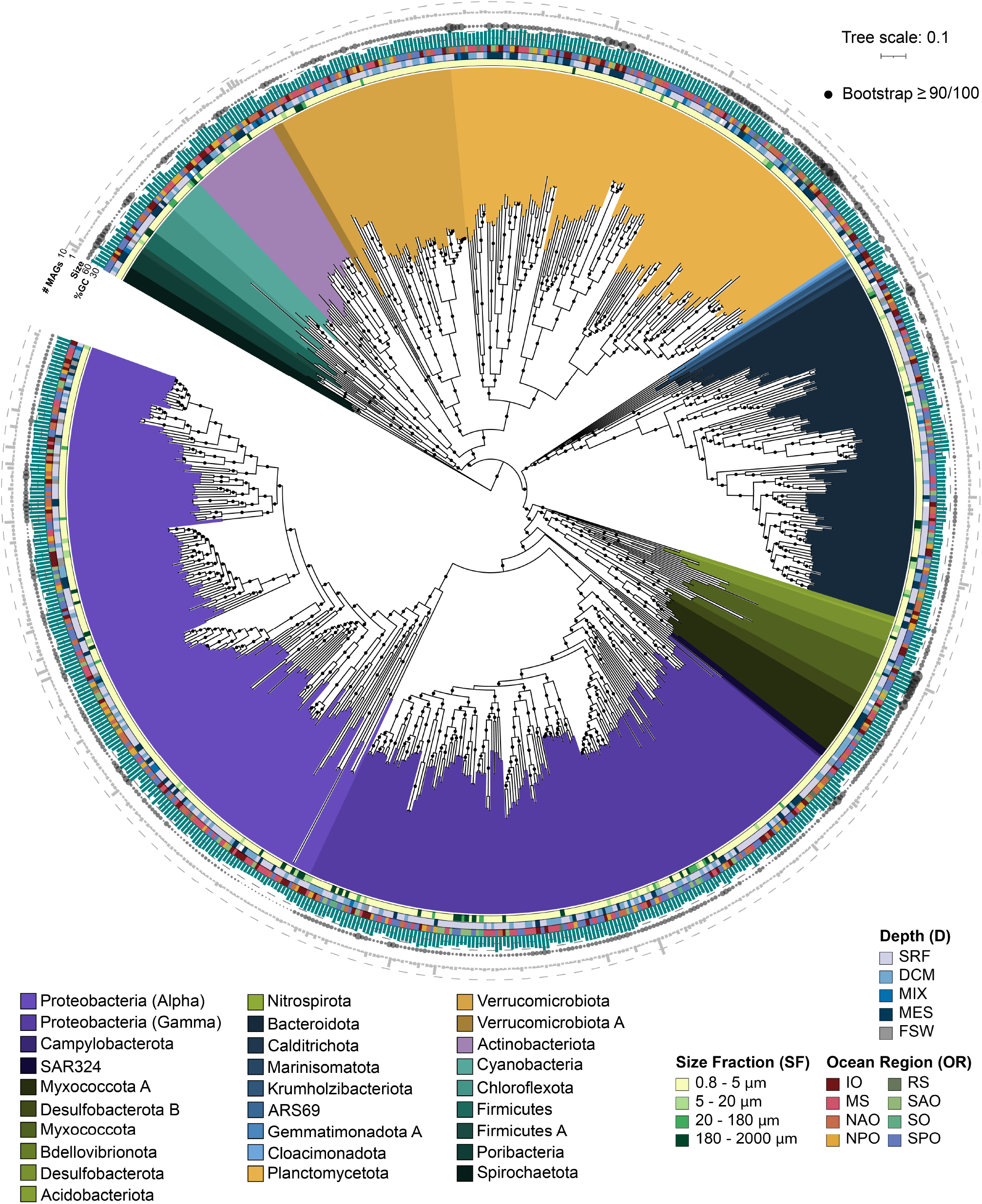
Diversity of the high-quality non-redundant bacterial TOPAZ MAGs. The approximately-maximum-likelihood phylogenetic tree was inferred form a concatenated protein alignment of 75 proteins using FastTree and GToTree workflow. The MAG names were omitted but the interactive version of the tree containing the MAG names can be accessed through iTOL (https://itol.embl.de/shared/halexand). Branches (nodes) are colored based on taxonomic annotations estimated by GTDBtk. The Ocean Region (OR), Size Fraction (SF), and Depth (D) of the co-assembly that a MAG was isolated from is color coded as colored bars. The GC (%) content is shown as a bar graph (in green), the genome size as bubble plot (the estimated size of the smallest genome included in this tree is 1.00Mbp and the largest is 13.24Mbp) and the number of MAGs in each genomic cluster (of 99 or higher %ANI) as a bar plot (in grey)

**Table 1:** Phylogenetic diversity and gain of various MAGs originating from *Tara* Oceans. Phylogenetic diversity and gain of prokaryotic MAGs was assessed for this study (TOPAZ), TOBG (*13*), UBA (*11*), and TARA (*12*) relative to each other as well as a “Neutral” tree comprised of relevant marine bacteria.

| Base tree | MAGs of interest | No. of MAGs | Phylogenetic diversity* | Phylogenetic gain° |
| --- | --- | --- | --- | --- |
| Neutral | TOPAZ (MQ, NR) | 1,571 | 59.9% | 42.8% |
| Neutral | TOPAZ (HQ, NR) | 634 | 41.6% | 25.8% |
| Neutral | TOBG | 1,974 | 61.3% | 46.7% |
| Neutral | UBA | 1,052 | 40.5% | 25.8% |
| Neutral | TARA | 722 | 44.4% | 31.0% |
| Neutral | TOBG + UBA + TARA | 3,750 | 66.6% | 51.8% |
| Neutral + *Tara* Oceans MAGs<sup>HQ</sup> | TOPAZ (HQ, NR) | 634 | 26.1% | 6.2% |
| Neutral + *Tara* Oceans MAGs<sup>MQ</sup> | TOPAZ (MQ, NR) | 1,572 | 44.7% | 14.4% |
| Neutral + *Tara* Oceans MAGs<sup>MQ</sup> | TOBG | 1,977 | 48.5% | 11.1% |
| Neutral + *Tara* Oceans MAGs<sup>MQ</sup> | UBA | 1,055 | 23.8% | 1.6% |
| Neutral + *Tara* Oceans MAGs<sup>MQ</sup> | TARA | 722 | 28.0% | 3.4% |

\* total branch length spanned by a set of taxa
° additional branch length contributed by a set of taxa
<sup>HQ</sup> includes Neutral, TOBG, UBA, TARA, and TOPAZ HQ, NR MAGs
<sup>MQ</sup> includes Neutral, TOBG, UBA, TARA, and TOPAZ MQ, NR MAGs

To confirm the hypothesis that the prokaryotic TOPAZ MAGs included particle-associated members, we examined the genomic features of several selected groups that were well-recovered here and in single-cell amplified genomic datasets (i.e., GORG) (*57*). To avoid potential biases related to completeness and contamination of the genomes, only the HQ-NR MAGs were compared to the GORG SAGs, and analyses were limited to groups with sufficient representation within both datasets (Bac-teriodota, Cyanobacteria, and Proteobacteria). For these well represented groups, the average GC% and estimated genome size of the TOPAZ MAGs were significantly higher than the ones typically reported in free-living marine bacteria (*58–60*) and those observed within the GORG dataset (*57*). TOPAZ MAGs were found to encode more tRNAs on average per genome than GORG (39.5 vs 30). Additionally, Carbohydrate-Active Enzymes (CAZy) and peptidases were enriched within the TOPAZ MAGs relative to GORG (Figure S22). Larger genomes have been considered diagnostic for a copiotrophic lifestyle in bacteria (*61*), since the more extended and flexible gene repertoire can facilitate substrate catabolism in organic rich niches such as particles. Genomes of copiotrophs are also commonly found to have higher copy numbers of genes associated with replication and protein biosynthesis such as tRNAs and rRNAs (*62*) which facilitate higher growth rates. In contrast, the streamlined genomes of SAR11 and other groups that have free-living oligotrophic lifestyles require fewer resources to maintain and replicate their genomes and have higher carbon-use efficiency (*63*). Similarly, G and C have higher energy cost of production and more limited intracellular availability compared to A and T (*60, 64*). The genomic trends observed support our findings that TOPAZ MAGs represent both particle associated and free-living microbes, and are relatively enriched for copiotrophic microbes.

### Environmental factors structure TOPAZ MAG co-occurrence

The co-retrieval of eukaryotic and prokaryotic MAGs from across the global ocean allows the unique opportunity to assess the biogeographical and ecological associations and potential co-occurrence of these organisms while also being able to infer likely function. To identify communities of associated organisms that co-occur across the surface ocean metagenomes, we performed a correlation clustering based on the abundances of the eukaryotic TOPAZ MAGS and the HQ-NR-TOPAZ MAGs (Figure 4a). We employed a modularity optimization algorithm to the correlation analysis (*67*) to identify distinct communities of co-occurring organisms. This approach identified five distinct communities (Figure 4 b). The communities were variably connected to each other, as defined by Equation (4), with highest connectedness between communities 1 and 2 and 4 and 5 (Figure 4 b; the maximum connectedness between 1 and 2 was 0.233 and the maximum connectedness between 4 and 5 was 0.448). Community 3 showed the lowest degree of connectivity within community members and to other communities (mean=0.108; remaining community mean=0.248), suggesting that members of this community co-occur less consistently across samples.

**Figure 4.**
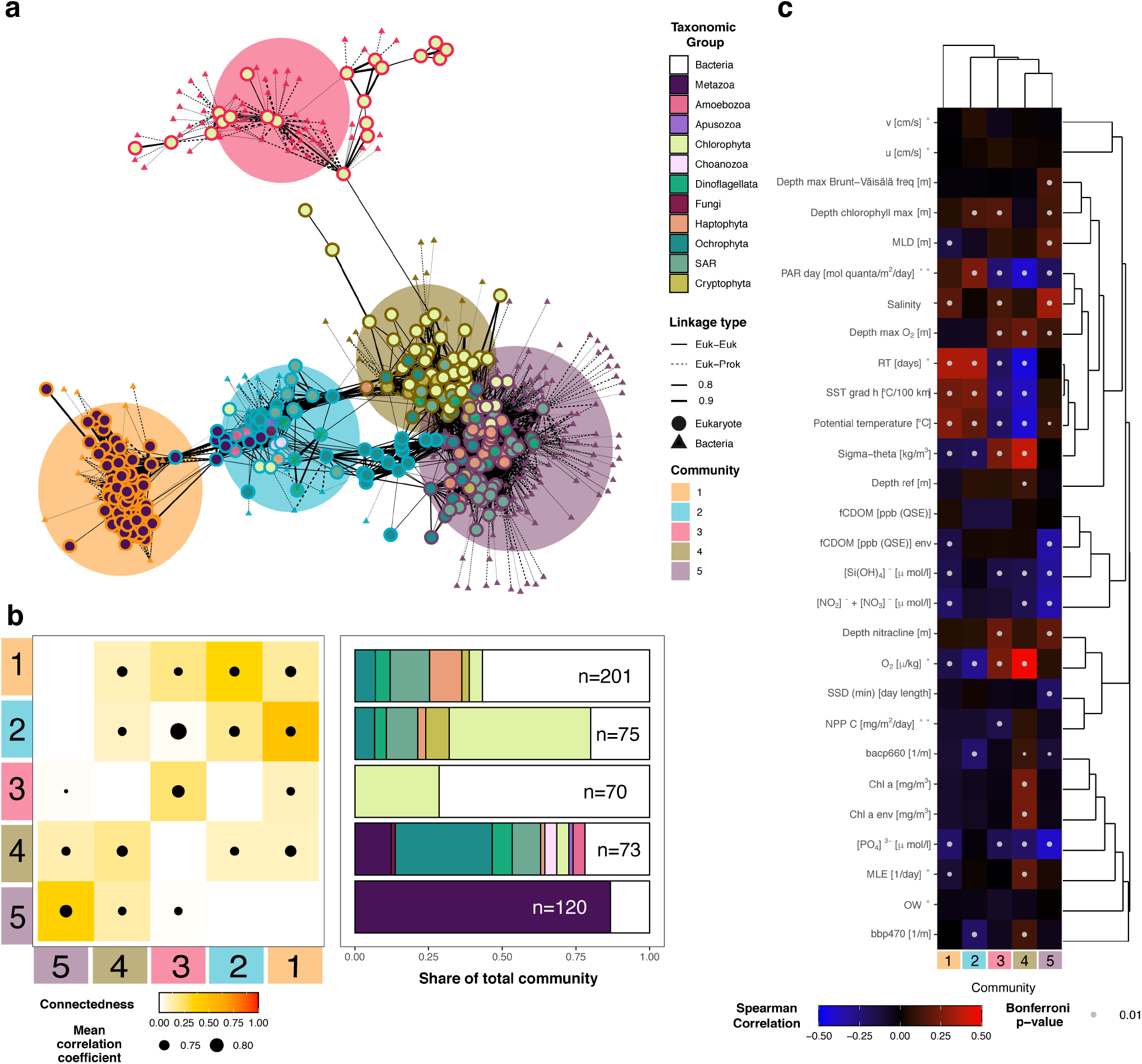
Distinct communities recovered from the TOPAZ MAGs. a) A network analysis performed on the metagenomic abundance of all recovered eukaryotic and prokaryotic TOPAZ MAGs based on Spearman Correlation analysis, identifying 5 distinct communities (see Materials and Methods). A force-directed layout of the seven communities is shown with eukaryotes (circles) and bacteria (triangles). Only linkages between eukaryotes are visualized. b) The connectedness and taxonomic composition of each community is depicted. Connectedness was calculated based on Equation (4). c) A Spearman correlation between the summed metagenomic abundance of each community and environmental parameters from the sampling (*65*), modeled mesoscale physical features based on d’Ovidio et al. (*66*) (indicated with *), and averaged remote sensing products (indicated with **). Significant Spearman correlations, those with a Bonferroni adjusted *p <* 0.01, are indicated with a dot on the heatmap.

The five communities that we identified based on metagenomic abundance correlations also significantly correlated with environmental factors, which consequently define the environmental niches where the communities were most abundant (Figure 4 c, Supplementary Table 13). Temperature was a primary factor defining the community correlations, significantly correlating with four of the five communities. Communities 2 and 4 correlated with colder temperatures (Figure 4 c). For Community 4, there was a significant positive correlation with chlorophyll (Chla: *ρ* = 0.236, *p* = 3.69*e* − 11), while we found negative correlations with “residence time” (*ρ* = − 0.438, *p* = 1.61*e* − 24), indicating a likely occurrence in newly formed eddies (according to the calculation by (*66*) as reported in the *Tara* Oceans metadata (*65*) as “residence time”). This aligns with the finding that Community 4 was typically found within colder, productive regions, and had higher metagenomic abundances in the Southern Ocean and the North Atlantic (Figure S24). The composition of Community 4 MAGs included Chlorophyta, Cryptophyta, Haptophyta, and Ochrophyta, the major groups containing primarily phototrophic eukaryotic microbes. 14 prokaryotic MAGs were also contained in this community, including both photosynthetic (Synechococcales) and non-photosynthetic lineages (e.g. Myxococcota and Planctomycetota). This guild of MAGs comprises likely photosynthesizers often found in cold, but not necessarily nutrient-rich, environments. Communities 1 and 3 correlated with warmer temperatures (Figure 4 c), which was attributed to their presence in longer-lived eddies (Figure 4 c; Community 1: *ρ* = 0.349, *p* = 1.19*e* − 14, Community 3: *ρ* = 0.345, *p* = 2.51*e* − 14). However, these two communities differed both in their association with nutrients and their taxonomic compositions. Community 1 was dominated by Metazoa and bacteria and correlated with oligotrophic conditions (nitrate and nitrite: *ρ* = − 0.218, *p* = 3.36*e* − 9, phosphate: *ρ* = − 0.218, *p* = 3.39*e* − 9, silica: *ρ* = − 0.157, *p* = 1.96*e* − 4), and was most abundant in the larger size fraction samples (20-2000 *µm*) (Figure S24). In contrast, Community 3 was largely comprised of phototrophic chlorophytes and bacteria and was not significantly correlated with nutrient conditions, and was most abundant in smaller size fraction samples (0.8-20 *µm*), particularly around the tropics (Figure S24). Community 5 was weakly associated with warmer water (*ρ* = 0.108,*p* = 9.10*e* − 2) and comprised by SAR and bacteria (Figure 4 b). Additionally, Community 5 was negatively correlated with nutrient concentrations (nitrate and nitrite: *ρ* = −0.348, *p* = 2.14*e* − 25, phosphate: *ρ* = −0.407, *p* = 7.36*e* − 36, silica: *ρ* = −0.310, *p* = 7.62*e* − 20), suggesting that this community thrives in oligotrophic regions.

While many of the communities recovered appeared to be driven largely by environmental forces, the genomic signatures of Community 1 members suggest that this community was comprised by MAGs from hosts and likely microbiome-associated microbes. Community 1 was comprised primarily of Metazoa, specifically Hexanauplia, and bacterial MAGs (Figure 4 b). Many of the bacterial MAGs in Community 1 had genes that suggest adaptations to microaerophic niches such as those which might be experienced when living in close host association (e.g. high affinity oxygen cytochromes, and reductases) (Figure S26). The bacterial MAGs in Community 1 could be broadly broken into two apparent functional types: those with larger genomes typical of copiotrophic bacteria and those with small genomes indicative potentially of reductive evolution. The first group was comprised of MAGs from family Saprospiraceae in phylum Bacteriodota (n=2, 3.0 Mbp average genome size), the order Opitutales in phylum Verrucomicrobiota (n=2, 3.4Mbp), and the family Vibrionaceae (n=2, 4.5 Mbp) in phylum Gammaproteobacteria (Figure S26). In addition to their relatively large size, the Saprospiraceae and Vibrionaceae MAGs were found to encode for genes involved in the hydrolysis and utilization of various complex carbon sources including chitin and other carbohydrates (Figure S26), such as those that might be shed or excreted by zooplankton such as copepods (*68*). By contrast, the second group of bacterial MAGs within Community 1 with smaller genomes, included MAGs from the Proteobacteria order Rickettsiales (n=3, 0.6-1.2 Mbp) and Gammaproteobacteria family Francisellaceae (n=1, 1.2 Mbp) Figure S26). The smaller genome sizes exhibited by these groups may be indicative of a genome streamlining which occurred with reductive evolution due to obligate or facultative symbiosis (*63*). Rickettsiales and Francisellaceae contain well-described obligate intracellular symbionts (*69–71*) and zoonotic pathogens (*70, 72*).

## Conclusion

Sequence datasets are revolutionizing how we form new hypotheses and explore environments on the planet. Here, we demonstrated a critical advance in the recovery of MAGs from environmentally-relevant eukaryotic organisms with EukHeist. The retrieval and study of MAGs to study the role of microorganisms in environmentally significant biogeochemical cycling is promising; however the current lack of eukaryotic reference genomes and transcriptomes complicates our ability to interpret the eukaryotic component of the microbial community. We recovered 988 total eukaryotic MAGs, 485 of which were deemed highly complete. Our findings demonstrate that specific branches of the eukaryotic tree were more likely to be resolved at the MAG-level due to their smaller genome size, distribution in the water column, and biological complexity. A substantial portion of the recovered eukaryotic MAGs were distinct from existing sequenced representatives, demonstrating that these large-scale surveys are a critical step towards characterizing less-resolved branches of the eukaryotic tree of life.

The continuing expansion of global-scale meta’omic surveys, such as BioGeoTraces (*73*), Bio-GO-SHIP (*74*), and the continuation of the Tara Oceans work highlights the importance of developing scalable and automated methods to enable more complete analysis of these data. Metagenomic pipelines that specifically integrate steps for handling eukaryotic biology, such as the EukHeist pipeline, are vital as eukaryotes are important members of microbial communities, ranging from the ocean, to soil (*75*) and human- (*76*) and animal-associated (*77*) environments. Additionally, we aim to contribute computational tools that can be integrated or customized, including EUKulele, EukMetaSanity, and eukrhythmic (*24, 26, 78*). The application of eukaryotic-sensitive methods such as EukHeist to other systems stands to greatly increase our understanding of the diversity and function of the “eukaryome”.

## Materials and Methods

### Data acquisition

The metagenomic and metatranscriptomic data corresponding to the size fractions dominated by eukaryotic organisms ranging from microbial eukaryotes and zooplankton (0.8 − 2000*µm*) as originally published by (*22*) were retrieved from the European Molecular Biology Laboratory-European Bioinformatics Institute (EMBL-EBI) under the accession numbers PRJEB4352 (large size fraction metagenomic data) and PRJEB6603 (large size fraction metatranscriptomic data) on November 20, 2018. Only samples with paired end reads (forward and reverse) were used in the subsequent analyses (Supplementary Table 1). After an initial sample-to-sample comparison with sourmash (sourmash compare -k 31 -scaled 10000) (*79*) (Figure S3), it was determined that samples largely clustered by depth and size fraction. Samples were grouped for co-assembly by size fraction (0.8 − 5*µm*, 5 − 20*µm*, 20 − 180*µm*, and 180 − 2000*µm*) as per (*22*), depth or sample type (surface (SRF), deep chlorophyll maximum (DCM), mesopelagic (MES), mixed surface sample (MIX), and filtered seawater (FSW)), and geographic location (Supplementary Table 1). In cases where a sample did not fall directly within one of the size classes, it was assigned to an existing size class based on the upper *µm* limit of the sample. This grouping resulted in the combination of 824 cleaned, paired FASTQ files samples into 94 distinct co-assembly groups, which were used downstream for co-assembly (Supplementary Table 1).

### EukHeist pipeline for metagenome assembly and binning

The metagenomic analysis, assembly, binning, and all associated quality control steps were carried out with a bioinformatic pipeline, EukHeist, that enables user-guided analysis of stand-alone metagenomic or paired metagenomic and metatranscriptomic sequence data. EukHeist is a streamlined and scalable pipeline currently based on the Snakemake workflow engine (*80*) that is configured to facilitate deployment on local HPC systems. Figure S2 outlines the structure and outputs of the existing EukHeist pipeline. EukHeist is designed to retrieve and identify both eukaryotic and prokaryotic MAGs from large, metagenomic and metatranscriptomic datasets (Figure S2). EukHeist takes input of sequence meta-data, user-specified assembly pairings (co-assembly groups), and raw sequence files, and returns MAGs that are characterized as either likely eukaryotic or prokaryotic.

Here, all raw sequences accessed from the EMBL-EBI were quality assessed with FastQC and MultiQC (*81*). Sequences were trimmed using Trimmomatic (v. 0.36; parameters: `ILLUMINACLIP: 2:30:7, LEADING:2, TRAILING:2, SLIDINGWINDOW:4:2, MINLEN:50`) (*82*). Passing mate paired reads were maintained for assembly and downstream analyses. Quality trimmed reads co-assembled based on assembly groups (Supplementary Table 1) with MEGAHIT (v1.1.3, parameters: k= 29, 39, 59, 79, 99, 119) (*83*). Basic assembly statistics were assessed for all co-assemblies with Quast (v. 5.0.2) (*84*) (Supplementary Table 1). Cleaned reads from assembly-group-associated metagenomic and metatranscriptomic samples were mapped back against the assemblies with bwa mem (v.0.7.17) (*85*). The bwa-derived abundances were summarized with MetaBat2 (v. 2.12.1) script jgi_summarize_bam_contig_depths (with default parameters). The output contig abundance tables were used along with tetranucleotide frequencies to associate contigs into putative genomic bins using MetaBat2 (v. 2.12.1) (*9*). The Snakemake profile used to conduct this analysis is available at https://www.github.com/alexanderlabwhoi/tara-euk-metag. A generalized version of the Snakemake pipeline (called EukHeist) that might be readily applied to other datasets is available at https://www.github.com/alexanderlabwhoi/EukHeist. MAGs here are subsequently named and referred to as **T**ara **O**ceans **P**article **A**ssociated MAGs (TOPAZ) and are individually named based on their assembly group (Supplementary Tables 2 and 3).

### Identification of putative Eukaryotic MAGs

The binning process described above recovered a total of 16,385 putative bins. These bins were screened to identify high completion eukaryotic and prokaryotic bins. All bins were first screened for length, assuming that eukaryotic bins would likely be greater than 2.5Mbp in size (modeled off of the size of the smallest known eukaryotic genome, ∼ 2.3Mbp *Microsporidian Encephalitozoon intestinalis* (*86*)). Bins larger than 2.5Mbp were screened for relative eukaryotic content using EukRep (*23*), a k-mer based strategy that estimates the likely domain-origin of metagenomic contigs. EukRep was used to classify the relative proportion of eukaryotic and prokaryotic content in each bin in a contig-by-contig manner. This approach identified 907 candidate eukaryotic bins that were greater than 2.5Mb in length and estimated to have more than 90% eukaryotic content by length. Protein coding domains were predicted in all 907 putative eukaryotic bins using EukMetaSanity (*24*).

### Protein prediction in Eukaryotic MAGs with EukMetaSanity

#### Taxonomy

The MMseqs2 v12.113e3 (*25,87,88*) `taxonomy` module (parameters: -`s 7 –min-seq-id 0.40 -c 0.3 –cov-mode 0`) was used to provide a first-pass taxonomic assignment of the input MAG for use in a downstream element of EukMetaSanity pipeline that requires an input NCBI taxon id or a taxonomic level (i.e. Order, Family, etc.). We created a custom database comprising both OrthoDB (*89*) and MMETSP (*43*) protein databases (OrthoDB-MMETSP) that integrates NCBI taxon ids. MMseqs2 was used to query each MAG against the OrthoDB-MMETSP database to identify a first-pass taxonomic assignment. The lowest common ancestor of top scoring hits was identified to provide taxonomic assignment to each candidate eukaryotic bin. The `taxonomyreport` module generates a taxon tree that includes the percent of MMseqs mappings that correspond to each taxonomic level. A taxonomic identifier and scientific name are selected to the strain level or when total mapping exceeds 8%, whichever comes first. The assigned NCBI taxon id is retained for downstream analyses.

#### Repeats identification

RepeatModeler (*90, 91*) was used to provide *ab initio* prediction of transposable elements, including short and long interspersed nuclear repeats, as well as other DNA transposons, small RNA, and satellite repeats. RepeatMasker (*92*) was then used to hard-mask these identified regions, as well as any Family-level (as identified above) repeats from the DFam 3.2 database (*93*). RepeatMasker commands `ProcessRepeats` (parameter: -`nolow`) and `rmOutToGff3` (parameter: `-nolow`) were used to output masked sequences (excluding low-complexity repeat DNA from the mask) as FASTA and gene-finding format (GFF3) files, respectively.

#### *Ab initio* prediction

GeneMark (*94*) was used to generate *ab initio* gene predictions with the repeat-masked eukaryotic candidate bin sequences output from the prior step. The GeneMark subprogram ProtHint attempts to use Order-level proteins from OrthoDB-MMETSP database to generate intron splice-site predictions for *ab initio* modeling using GeneMark EP (*95*). If ProtHint fails to generate predictions, then GeneMark will default to ES mode. Due to the fragmented nature of metagenomic assemblies, the prediction parameter stringency was drastically reduced relative to what is recommended for draft genome projects (parameters: `–min_contig 500 –min_contig_in_predict 500 –min_gene_in_predict 100`). These parameters can be easily modified within the EukMetaSanity config file. GeneMark outputs predictions of protein coding sequences (CDS) and exon/intron structure as GFF3 files.

#### Integrating protein evidence

MetaEuk (*96*) was used to directly map the repeat-masked eukaryotic candidate bins sequences against proteins from the MMETSP (*43, 97*) and eukaryotes included in the OrthoDB v10 dataset (*89*), hereafter referred to as the OrthoDB-MMETSP database. MetaEuk easy-predict (parameters: `–min-length 30 –metaeuk-eval 0.0001 -s 7 –cov-mode 0 –c 0.3 -e 100 –max-overlap 0`) used Order-level proteins to identify putative CDS and exon/intron structure. MetaEuk encodes this output as headers in FASTA sequences that are then parsed into GFF3 files.

#### Merging final results

GFF3 output from the previous *ab initio* and MetaEuk protein evidence steps were input into Gffread (*98*) (parameters: `-G –merge`) to localize predictions from both lines of evidence into a single GFF3 output file. Each locus was then merged together using a Python (*99*) script and the BioPython API (*100*) within EukMetaSanity. The set of *ab initio* generated exons in each locus is used as a prediction of the underlying exon/intron structure of the gene locus to which it is assigned. If there are any protein-evidence-generated exons present at the same locus, and if the total numbers of exons predicted by each line of evidence have ≥ 70% agreement, *ab initio* generated exons lacking a corresponding protein-evidence-generated exon are removed (the first and last exon(s) of a locus are not removed). Conversely, any protein-evidence-generated exon present that lacks a corresponding *ab initio* generated exon is added to the predicted exon/intron structure. The final gene structure for each locus is then processed into GFF3 and FASTA format.

### Functional and taxonomic annotation of eukaryotic MAGs

Predicted proteins from EukMetaSanity were annotated for function against protein families in Pfam with PfamScan (*101*) and KEGG using kofamscan (*102, 103*) (Supplementary Tables 7 and 8). The relative completeness and contamination of each putative Eukaryotic MAG was assessed based on protein content using BUSCO v 4.0.5 against the eukaryota_odb10 gene set using default parameters (*27*) and EukCC v 0.2 using the EukCC database (created 22 October 2019 (*104*)). Annotation and completeness assessment were carried out using a EukHeist-Annotate (https://www.github.com/halexand/EukHeist-annotate). EukCC (*104*) was also used to calculate MAG completeness and contamination. The average completeness across groups increased in all cases with EukCC except for metazoans, which on average had a lower estimated completeness (Figure S10).

The taxonomic affiliation of the high-and low-completion bins was estimated using MMSeqs taxon-omy through EukMetaSanity and EUKulele (*26*), an annotation tool that takes a protein-consensus approach, leveraging a Last Common Ancestor (LCA) estimation of protein taxonomy, as well as MMSeqs2 taxonomy module (*25, 87, 88*). Taxonomic level estimation in EUKulele was assessed based on e-value derived best-hits, where percent id was used as a means of assessing taxonomic level, with the following cutoffs: species, >95%; genus, 95-80%; family, 80-65%; order, 65-50%; class, 50-30% modeled off of Carradec et al. (2018). All MAGs were searched against the MarMet-Zoan combining the MarRef, MMETSP, and metazoan orthoDB databases (*43, 89, 97, 105*). MAGs with taxonomy assignment that did not resolve beyond SAR (Stramenopile, Alveolate, Rhizaria) are classified as SAR. This database is available for download through EUKulele.

### Phylogeny of eukaryotic MAGs

A total of 49 BUSCO proteins were found to be present across 80% or more of the highly complete eukaryotic TOPAZ MAGs and were selected for the construction of the tree. Amino acid sequences from all genomes and transcriptomes of interest were collected and aligned individually using mafft (v7.471) (parameters: `–thread -8 –auto`) (*106*). Individual protein alignments were trimmed to remove sections of the alignment that were poorly aligned with trimAl (v1.4.rev15) (parameters: -automated1) (*107*). Protein sequences were then concatenated and trimmed again with trimAl (parameters: -automated1). A final tree was then constructed using RAxML (v 8.2.12; parameters: `raxmlHPC-PTHREADS-SSE3 -T 16 -f a -m PROTGAMMAJTT -N 100 -p 42 -x 42`) (*108*). The amino acid alignment and construction was controlled with a Snakemake workflow: https://github.com/halexand/BUSCO-MAG-Phylogeny/. Trees were visualized and finalized with iTOL (*109*).

### Prokaryotic MAG assessment and analysis

The 15,478 bins that were not identified as putative eukaryotic bins based on length and EukRep metrics were screened to identify quality prokaryotic bins. The quality and phylogenetic-association of these bins was assessed with a modified version of MAGpy (*110*), which was altered to include taxonomic annotation with GTDB-TK v.0.3.2 (*111*). Bins were assessed based on single copy ortholog content with CheckM v (*112*) to identify 2 different bin quality sets: 1) high-quality (HQ) prokaryotic bins (>90% completeness, <5% contamination), and 2) medium-quality (MQ) prokaryotic bins (90-75% completeness, <10% contamination). A total of 4022 prokaryotic MAGs met the above criteria. A final set of 2,407 non-redundant (NR) HQ-MQ MAGs were identified using dRep v2.6.2 (*113*), which performs pairwise genome comparisons in two steps. First, a rapid primary algorithm, Mash v1.1.1 (*114*) is applied. Genomes with Mash values equivalent to 90% Average Nucleotide Identity (ANI) or higher were then compared with MUMmer v3.23 (*115*). Genomes with ANI ≥ 99% were considered to belong to the same cluster. The best representative MAGs were selected based on the dRep default scoring equation (*113*). Out of the final set of 2,407 NR MAGs, 716 were HQ. The same pipeline was used to determine the HQ and MQ NR MAGs reconstructed from the *Tara* Oceans metagenomes in previous studies (*11–13*).

### Phylogeny of bacterial non-redundant high-quality MAGs

Only 5 out of the 716 HQ NR MAGs were found to belong to Archaea, thus only bacterial MAGs were used for the construction of the phylogenetic tree with GToTree v.1.4.10 (*116*) and the gene set (HMM file) for Bacteria (74 targets). GToTree pipeline uses Prodigal v2.6.3 (*117*) to retrieve the coding sequences in the genomes, and HMMER3 v3.2.1 (*118*) to identify the target genes based on the provided HMM file. MUSCLE v3.8 (*119*) was then used for the gene alignments, and Trimal v1.4 (*107*) for trimming. The concatenated aligned is used for the tree constructions using FastTree v2.1 (*120*). Three genomes were excluded from the analysis due to having too few of the target genes. The tree was visualized using the Interactive Tree of Life (iToL) (*109*).

### Prokaryote MAG phylogeny comparison

A set of 8,644 microbial genomes were collected from the MarDB database (*105*)(accessed 31 May 2018) encompassing the publicly available marine microbial genomes. Genomes were assessed using CheckM v1.1.1 (*112*)(parameters: `lineage_wf`) and genomes estimated to be <70% complete or >10% contamination were discarded. The remaining genomes (n = 5,878) were assessed using CompareM v0.0.23 (parameters: `aai_wf;` https://github.com/dparks1134/CompareM) and near identical genomes were identified using a cutoff of ≥ 95% average amino acid identity (AAI) with ≥ 85% orthologous fraction (determined as one standard deviation from the average orthologous fraction for genomes with 97 − 100% AAI). Based on CheckM quality, the genome with the highest completion and/or lowest contamination were retained. From the remaining genomes (n = 3,843), all MAGs derived from the *Tara* Oceans dataset, specifically from Tully et al. (*13*) and Parks et al. (*11*), were removed. The remaining genomes (n = 2,275) would be used to form the base of a phylogenetic tree representing the available genome diversity prior to the release of previous *Tara* Oceans related MAG datasets (*11–13*), termed the “neutral” component of subsequent phylogenetic trees.

For the comparisons, phylogenetic trees were constructed using GToTree v1.4.7 (*116*) (default parameters; 25 Bacteria_and_Archaea markers). Any genome added to a tree that did not meet the default 50% marker presence requirement was excluded from that tree. Five iterations of phylogenetic trees were constructed using the neutral genomes paired with each *Tara* Oceans MAG dataset, the high-quality TOPAZ prokaryote MAGs, and the medium-quality TOPAZ prokaryote MAGs, individually, and two larger trees were constructed containing all neutral genomes and *Tara* Oceans MAGs, with additions of either high-or medium-quality TOPAZ MAGs. Phylogenetic trees were assessed using genometreetk (parameter: pd; https://github.com/dparks1134/GenomeTreeTk) to determine the phylogenetic diversity (i.e., the total branch length traversed by a set of leaves) and phylogenetic gain (i.e., the additional branch length added by a set of leaves) (*11*) for each set of MAGs compared against the neutral genomes and for the TOPAZ prokaryote MAGs compared against the neutral genomes and the other *Tara* Oceans MAGs.

### MAG abundance profiling

Raw reads from all metagenomic and metatranscriptomic samples were mapped against the eukaryotic and prokaryotic TOPAZ MAGs to estimate relative abundances with CoverM (v. 0.5.0; parameters: –min-read-percent-identity 0.95 –min-read-aligned-percent 0.75 –dereplicate –dereplication-ani 99 –dereplication-aligned-fraction 50 –dereplication-quality-formula dRep –output-format dense –min-covered-fraction 0 –contig-end-exclusion 75 –trim-min 0.05 –trim-max 0.95 –proper-pairs-only; https://github.com/wwood/CoverM). The total number of reads mapped to each MAG was then used to calculate Reads Per Kilobase Million (*RPKM*), where for some MAG, *i* : *RPKM*_*i*_ = *X*_*i*_*/l*_*i*_*N*10^9^, with *X* = total number of reads recruiting to a MAG, *l* = length of MAG in Kb, *N* = total number of trimmed reads mapping to a sample in millions. We also calculated counts per million (CPM), a normalization of the RPKM to the sum of all RPKMs in a sample. CPM, a modification of transcripts per million (TPM) was first proposed by (*121*) as an alternative to RPKM that reduces statistical bias. The metric has since been applied to metagenomics data, sometimes called GPM (genes per million) (*122*). We chose not to more stringently cluster MAGs on the basis of genome content due to the documented utility of preserving population-specific genes (*123*); we show a comparison of the CoverM-based dereplication approach using fastANI to the dnadiff function of the MUMmer paper in Figure S25.

### Nutritional modelling

To predict the trophic mode of the high quality TOPAZ eukaryotic MAGs (n=485), a Random Forest model (*124*) was constructed and calibrated using the ranger (*125*) and tuneRanger packages in R (*126*), respectively. The model was trained using KEGG Orthology (KO) annotations (*102*) from a manually-curated reference trophic mode transcriptomic dataset consisting of the MMETSP (*43*) and EukProt (*127*) (Supplementary Table 5). 644 of the transcriptomes in this reference dataset came from the MMETSP (*43*), after 22 transcriptomes were removed due to low coverage of KEGG and Pfam annotations (*101*). The remaining 266 came from the EukProt database (*127*), after 162 were removed due having fewer than 500 present KOs. Nutritional strategy (phototrophy, heterotrophy, or mixotrophy) was assessed for each reference transcriptome individually based on the literature, 25% of the combined reference transcriptomes were excluded from model training as testing data (Figure S16).

A subset of KEGG Orthologs (KOs) that were predictive for trophic mode classification was determined computationally with the vita variable selection package in R (*128*) (Supplementary Table 6), which has been tested and justified for this use case (*129*). This process was carried out by the algorithm without regard to the predicted function of the KOs, but we found that many of these KOs were implicated in carbohydrate and energy metabolism, with preference for those KOs that differ strongly between heterotrophs and phototrophs (particularly for energy metabolism; Figure S18). The model was built using the selected KOs (*n* = 1787 of a total 21585 KOs) with the 75% of the combined database assigned as training data.

Additionally, we developed a secondary metric for assessing the extent of heterotrophy of a transcriptome or MAG. As opposed to the trinary classification scheme of the Random Forest model, this approach quantifies the extent that the MAG aligns with heterotrophic, phototrophic, or mixotrophic references by assigning a composite score. We calculated the likelihood of vita selected KOs used in the Random Forest model above to be present within heterotrophic, phototrophic, or mixotrophic reference transcriptomes. Three scores (*h, p, m*), one corresponding to each trophic mode, were hence calculated for each vita-selected KO (*k*) (*n* = 1787) (Supplementary Table 6). In Equation (1), *K* is the number of references the KO was present in for each trophic mode category, while *n* is the total number of references available for each trophic mode category.

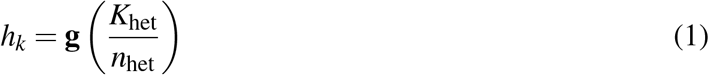

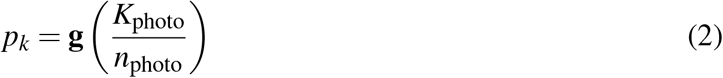

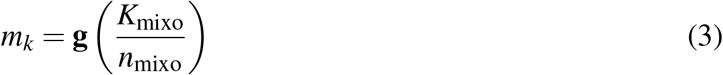

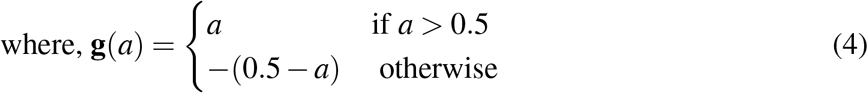

If a given KO occurred in fewer than 50% of the reference transcriptomes for a trophic mode, it was considered not to be characteristic of that trophic mode and as such the score, which we represent as the variable *a*, the ratio of the present KOs to the total for the subset of transcriptomes annotated some trophic mode (Equation (4)), was transformed (− (0.5 − *a*), if *a <* 0.5), to reflect the absence without valuing absence over presence. In the test transcriptome dataset, the ratio-transformed scores were negated when a given KO was absent from the transcriptome. For instance, if a KO was absent from 90% of reference transcriptomes assigned to heterotrophy (*a* = 0.1), and absent in the MAG or transcriptome being evaluated, it would receive a score of *h*_*k*_ = − 1 ∗ ((0.5 − 0.1)) = 0.4 (Equation (1)) for that KO. This reflects that the absence of the KO in the evaluated MAG or transcriptome aligned well with the high probability that the KO was absent among the reference transcriptomes.

The scores for all KOs selected by vita were then used to scale the presence/absence patterns observed across transcriptomes and MAGs. Thus, for each transcriptome or MAG a single score was calculated for each trophic mode heterotrophy (*H*), phototrophy (*P*), and mixotrophy (*M*) for all KOs present within the transcriptome or MAG (*K*):

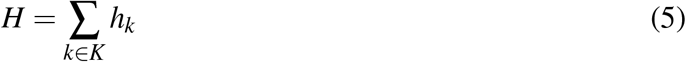

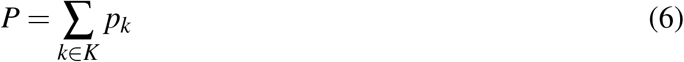

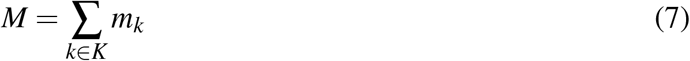

These calculated values can then be aggregated to a composite heterotrophy score (*H*_*ind*_) (Supplementary Table 9). The score was computed as follows:

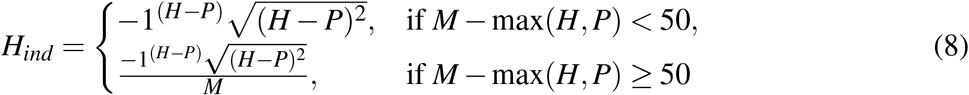

### Network Analysis

To identify co-occurring MAGs across the stations surveyed by *Tara* Oceans, the CPM abundance of each highly-complete eukaryotic MAG (*>* 30% BUSCO completeness) and each non-redundant, highly complete bacterial MAG was assessed at each station at all available depths and size fractions as described above. CPM was used because of the power of this metric for comparing samples directly: the sum of all CPM values per sample will be the same, as sequencing depth is accounted for after gene length. This makes it easier to compare the abundances of MAGs originally recovered from different sites (*122*). A Spearman correlation matrix was generated to identify monotonic relationships between MAGs. Correlations were filtered based first on p-value, using the Šidák correction (*130*), a slightly less stringent metric than the Bonferroni correction. The Šidák correlation adjusts for multiple comparisons and is given by *p <* 1 − (1 − *α*)^1*/n*^, where *n* is the total number of comparisons, and *α* is the significance value, in this case 0.05. We considered only those correlations within the 90th percentile of CPM correlations, thus correlations with absolute value less than 0.504 were removed from the analysis. Subsequently, we further filtered interactions to those with coefficient of correlation *>* 0.70 for the construction of the network diagram. Because it was expected for several of the eukaryotic MAGs to be closely related (based on ANI), the relationships in the network were further filtered to exclude interactions between MAGs of exceedingly high similarity (having both 99% ANI similarity and *>* 0.70 coefficient of correlation in the network analysis) (Supplementary Table 12). ANI-based group members tended to have identical taxonomic classifications: only 2 of 94 clusters had different classifications at the order level per EUKulele (**??**).

We generated a network from this reduced set of labeled interactions (cut off at *>* 0.70 coefficient of correlation, focusing on interactions between eukaryotes and prokaryotes or eukaryotes and eukaryotes, and using ANI-based clusters instead of MAG names when applicable) using `igraph` (*131, 132*) (Supplementary Table 11). Communities of highly associated MAGs were identified using a modularity optimization algorithm introduced in (*67*) and implemented in `igraph` (*131*).

We assessed the connectedness within and between communities by calculating a connectedness metric as follows. For the connectedness within a community (one community to itself), we identified the number of “dense” connections by counting up the total number of links found between community members, regardless of how many times the particular MAG had been connected to its own community, and divided that number by the total possible “dense”, meaning the number of connections which would exist if all community members were connected to all other community members. Between different communities, we defined connectedness by qualifying that a “connection” is made the first time each MAG from a given community is linked to another community, and calculated this quantity by dividing the number of realized links between community members by the maximum total size of the two involved communities (Figure 4 b; Equation (9) - Equation (11)).

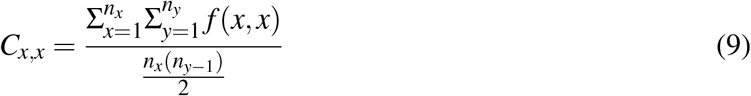

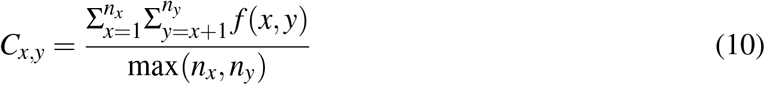

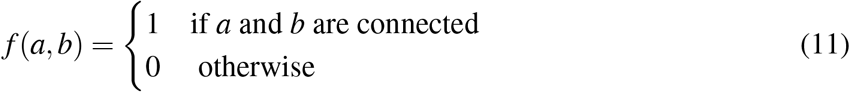

We calculated Spearman correlation coefficients for the relationship between the abundance of communities between stations and several environmental parameters of interest from the *Tara* Oceans metadata (*65, 133*) (Figure 4). We considered the measured physical and chemical parameters, the modeled mesoscale physical oceanographic parameters, and averaged remote sensing products (*65, 66, 133*). We adjusted the p-value of these comparisons using a Bonferroni adjustment within the statistics package in R (*132*).

### Data and code availability

The eukaryotic and prokaryotic TOPAZ MAGs and Supplementary Tables 1-13 are available through the Open Science Framework (OSF) at https://osf.io/gm564/ with the DOI: 10.17605/OSF.IO/GM564. EukHeist, which was used to recover the reported TOPAZ MAGs can be found at https://github.com/AlexanderLabWHOI/EukHeist and EukMetaSanity which was used for protein prediction in eukaryotic MAGs can be found at https://github.com/cjneely10/EukMetaSanity. Code used to generate the figures in this paper can be found at https://github.com/AlexanderLabWHOI/2021-TOPAZ-MAG-Figures. An interactive visualizer for the TOPAZ eukaryotic MAGs is available at https://share.streamlit.io/cjneely10/tara-analysis/main/TARAVisualize/main.py with source code at https://github.com/cjneely10/TARA-Analysis.

## Acknowledgements

This research would not have been possible without the community-driven efforts to provide open and freely available data by the *Tara* Oceans Consortium. HA and SKH conceived of and designed the study. HA carried out the assembly and binning. HA, AIK, SKH, MP, TR, and CJN, and BJT analyzed the data. HA and SKH wrote the manuscript with input from all authors. All authors edited and commented on the manuscript. This research was supported by a National Science Foundation grant (OCE-1948025) to HA and a WHOI Independent Research and Development award to HA. SKH was supported through a Postdoctoral Fellowship (OCE-0939564) provided through the NSF Center for Dark Energy Biosphere Investigations and through an NSF grant (OCE-1947776). AIK was supported by the Computational Science Graduate Fellowship (DOE; DE-SC0020347). MP was supported by a NSF grant (OCE-1924492) and a WHOI Independent Research and Development award. BJT was supported by the Center for Dark Energy Biosphere Investigations (C-DEBI) through NSF-OCE-0939654. The authors declare no conflicts of interest.

## References

1. D. A. Caron, P. D. Countway, A. C. Jones, D. Y. Kim, A. Schnetzer, Annual Review of Marine Science 4, 467 (2011).

2. A. Mitra, et al., Biogeosciences 11, 995 (2014).

3. D. A. Caron, et al., Nature Reviews Microbiology 15, 6 (2017).

4. S. L. Strom, Science 320, 1043 (2008).

5. D. A. Caron, P. D. Countway, Aquatic Microbial Ecology 57, 227 (2009).

6. P. J. Keeling, d. J. Campo, Current Biology 27, R541 (2017).

7. J. Alneberg, et al., Nature Methods 11, 1144 (2014).

8. Y.-W. Wu, Y.-H. Tang, S. G. Tringe, B. A. Simmons, S. W. Singer, Microbiome 2, 26 (2014).

9. D. D. Kang, et al., PeerJ 7, e7359 (2019).

10. E. D. Graham, J. F. Heidelberg, B. J. Tully, PeerJ 5, e3035 (2017). NULL.

11. D. H. Parks, et al., Nature Microbiology 2, 1533 (2017).

12. T. O. Delmont, et al., Nature Microbiology 3, 804 (2018).

13. B. J. Tully, E. D. Graham, J. F. Heidelberg, Scientific Data 5, 170203 (2018).

14. A. Almeida, et al., Nature 568, 499 (2019).

15. C. Rinke, et al., ISME Journal 13, 663 (2019).

16. B. J. Tully, Nature Communications 10, 1 (2019).

17. T. O. Delmont, et al., Cell Genomics p. 100123 (2022).

18. A. Duncan, et al., Microbiome 10, 1 (2022).

19. R. Massana, D. López-Escardó, Cell Genomics 2, 100130 (2022).

20. W. Zhang, et al., PLoS ONE 6 (2011).

21. M. R. Olm, et al., Microbiome 7, 26 (2019).

22. Q. Carradec, et al., Nature Communications 9, 373 (2018).

23. P. T. West, A. J. Probst, V. I. Grigoriev, B. C. Thomas, J. F. Banfield, Genome research 28, 569 (2018).

24. C. J. Neely, S. K. Hu, H. Alexander, B. J. Tully, bioRxiv (2021).

25. M. Steinegger, J. Söding, Nature Communications 9, 1 (2018).

26. A. I. Krinos, S. K. Hu, N. R. Cohen, H. Alexander, Journal of Open Source Software (2021).

27. F. A. Simao, R. M. Waterhouse, P. Ioannidis, V. E. Kriventseva, E. M. Zdobnov, Bioinformatics 31, 3210 (2015).

28. Y. Hou, S. Lin, PLoS ONE 4, e6978 (2009).

29. B. A. Read, et al., Nature 499, 209 (2013).

30. C. Bowler, et al., Nature 456, 239 (2008).

31. I. H. G. S. Consortium, Nature 431, 931 (2004).

32. F. Burki, A. J. Roger, M. W. Brown, A. G. Simpson, Trends in Ecology & Evolution 35, 43 (2020).

33. T. S. Jørgensen, et al., G3 Genes|Genomes|Genetics 9, 1295 (2019).

34. S. E. Morales, A. Biswas, G. J. Herndl, F. Baltar, Front. Mar. Sci. 6 (2019).

35. M. C. Pernice, et al., ISME Journal 10, 945 (2015).

36. V. P. Edgcomb, D. Beaudoin, R. Gast, J. F. Biddle, A. Teske, Environmental Microbiology 13, 172 (2010).

37. T. K. Mohanta, H. Bae, Biol Proced Online 17 (2015).

38. M. C. Pernice, et al., ISME Journal 9, 782 (2014).

39. A. Z. Worden, et al., Science 347, 1257594 (2015).

40. C. de Vargas, et al., Science 348, 1261605 (2015).

41. E. B. Sherr, B. F. Sherr, Antonie van Leeuwenhoek 81, 293 (2002).

42. D. K. Stoecker, P. J. Hansen, D. A. Caron, A. Mitra, Annu. Rev. Mar. Sci. 9, 311 (2017).

43. P. J. Keeling, et al., PLOS Biology 12, 1 (2014).

44. Z. Liu, V. Campbell, K. B. Heidelberg, D. A. Caron, FEMS Microbiology Ecology 92, fiw106 (2016).

45. H. Alexander, et al., Proceedings of the National Academy of Sciences 112, E5972 (2015).

46. S. K. Hu, et al., Environmental Microbiology 20, 2865 (2018).

47. W. Gong, et al., ISME Journal 11, 439 (2016).

48. A. Labarre, A. Obiol, S. Wilken, I. Forn, R. Massana, Limnol Oceanogr 65 (2020).

49. M. A. Shipp, et al., Nature Medicine 8, 68 (2002).

50. A. Bashiri, M. Ghazisaeedi, R. Safdari, L. Shahmoradi, H. Ehtesham, Iranian Journal of Public Health 46, 165 (2017).

51. A. A. Tabl, A. Alkhateeb, W. ElMaraghy, L. Rueda, A. Ngom, Frontiers in Genetics 10, 256 (2019).

52. E. H. Mahood, L. H. Kruse, G. D. Moghe, Applications in Plant Sciences 8, e11376 (2020).

53. B. S. Lambert, et al., Proceedings of the National Academy of Sciences 119, e2100916119 (2022).

54. J. A. Burns, A. A. Pittis, E. Kim, Nature Ecology & Evolution 2, 697 (2018).

55. V. Jimenez, J. A. Burns, F. L. Gall, F. Not, D. Vaulot, J. Phycol. 57, 435 (2021).

56. K. J. Flynn, et al., Journal of Plankton Research 41, 375 (2019).

57. M. G. Pachiadaki, et al., Cell 179, 1623 (2019).

58. A. Dufresne, L. Garczarek, F. Partensky, Genome Biol 6, R14 (2005).

59. B. K. Swan, et al., Proceedings of the National Academy of Sciences 110, 11463 (2013).

60. H. Luo, L. R. Thompson, U. Stingl, A. L. Hughes, Mol Biol Evol 32, 2738 (2015).

61. J. G. Okie, et al., eLife 9 (2020).

62. E. P. Rocha, Genome Research 14, 2279 (2004).

63. S. J. Giovannoni, J. C. Thrash, B. Temperton, ISME Journal 8, 1553 (2014).

64. C. M. Moore, et al., Nature Geosci 6, 701 (2013).

65. C. Tara Oceans Consortium, P. Tara Oceans Expedition, Environmental context of all samples from the Tara Oceans Expedition (2009-2013), about water column features (PANGAEA, 2016). In: Tara Oceans Consortium, C; Tara Oceans Expedition, P (2016): Registry of all samples from the Tara Oceans Expedition (2009-2013). PANGAEA, https://doi.org/10.1594/PANGAEA.859953.

66. F. d’Ovidio, S. D. Monte, S. Alvain, Y. Dandonneau, M. Levy, Proceedings of the National Academy of Sciences 107, 18366 (2010).

67. V. D. Blondel, J.-L. Guillaume, R. Lambiotte, E. Lefebvre, Journal of Statistical Mechanics: Theory and Experiments 2008, P10008 (2008).

68. D. D. Corte, et al., Environ Microbiol 20, 492 (2017).

69. D. Santos-Garcia, et al., Genome Biology and Evolution 6, 1013 (2014).

70. A. C. Darby, N.-H. Cho, H.-H. Fuxelius, J. Westberg, S. G. Andersson, Trends in Genetics 23, 511 (2007).

71. D. Li, J. Fang, B. Wen, X. Wu, Aquaculture 539, 736565 (2021).

72. J. Celli, T. C. Zahrt, Cold Spring Harbor Perspectives in Medicine 3, a010314 (2013).

73. S. J. Biller, et al., Sci Data 5 (2018).

74. L. J. Ustick, et al., Science 372, 287 (2021).

75. J. Bailly, et al., The ISME journal 1, 632 (2007).

76. J. Lukeš, C. R. Stensvold, K. Jirků-Pomajbíková, L. W. Parfrey, PLoS Pathogens 11, e1005039 (2015).

77. J. Campo, D. Bass, P. J. Keeling, Funct Ecol 34, 2045 (2019).

78. A. I. Krinos, N. R. Cohen, M. J. Follows, H. Alexander, bioRxiv (2022).

79. C. T. Brown, L. Irber, JOSS 1, 27 (2016).

80. J. Koster, S. Rahmann, Bioinformatics 28, 2520 (2012).

81. S. Andrews, Fastqc: A quality control tool for high throughput sequence data. (2010). [Online; accessed 2014-03-31].

82. A. M. Bolger, M. Lohse, B. Usadel, Bioinformatics 30, 2114 (2014).

83. D. Li, C.-M. Liu, R. Luo, K. Sadakane, T.-W. Lam, Bioinformatics 31, 1674 (2015).

84. A. Gurevich, V. Saveliev, N. Vyahhi, G. Tesler, Bioinformatics 29, 1072 (2013).

85. H. Li, R. Durbin, Bioinformatics 26, 589 (2010).

86. N. Corradi, J. F. Pombert, L. Farinelli, E. S. Didier, P. J. Keeling, Nature Communications 1 (2010).

87. M. Steinegger, J. Söding, MMseqs2 enables sensitive protein sequence searching for the analysis of massive data sets (2017).

88. M. Mirdita, M. Steinegger, J. Söding, Bioinformatics 35, 2856 (2019).

89. E. V. Kriventseva, et al., Nucleic Acids Research 47, D807 (2018).

90. J. M. Flynn, et al., Proceedings of the National Academy of Sciences of the United States of America 117, 9451 (2020).

91. A. Smit, R. Hubley, Repearmodeler open-1.0, http://www.repeatmasker.org (2008-2015).

92. A. Smit, R. Hubley, P. Green, Repeatmasker open-4.0, http://www.repeatmasker.org (2013-2015).

93. J. M. Flynn, et al., Proceedings of the National Academy of Sciences of the United States of America 117, 9451 (2020).

94. A. Lomsadze, V. Ter-Hovhannisyan, Y. O. Chernoff, M. Borodovsky, Nucleic Acids Research 33, 6494 (2005).

95. T. Bruna, A. Lomsadze, M. Borodovsky, NAR Genomics and Bioinformatics 2 (2020).

96. E. Levy Karin, M. Mirdita, J. Söding, Microbiome 8, 48 (2020).

97. L. K. Johnson, H. Alexander, C. T. Brown, GigaScience (2018).

98. M. Pertea, G. Pertea, F1000Research 9, 304 (2020).

99. P. S. Foundation, Python language reference, version 3.6, http://www.python.org.

100. P. J. A. Cock, et al., Bioinformatics 25, 1422 (2009).

101. R. D. Finn, et al., Nucleic Acids Research 42, D222 (2014).

102. M. Kanehisa, Protein Science 28, 1947 (2019).

103. T. Aramaki, et al., Bioinformatics 36, 2251 (2019).

104. P. Saary, A. L. Mitchell, R. D. Finn, Genome Biology 21 (2020).

105. T. Klemetsen, et al., Nucleic Acids Research 46, D692 (2017).

106. K. Katoh, D. M. Standley, Molecular Biology and Evolution 30, 772 (2013).

107. S. Capella-Gutierrez, J. M. Silla-Martinez, T. Gabaldon, Bioinformatics 25, 1972 (2009).

108. A. Stamatakis, Bioinformatics 30, 1312 (2014).

109. I. Letunic, P. Bork, Nucleic acids research 44, W242 (2016).

110. R. D. Stewart, M. D. Auffret, T. J. Snelling, R. Roehe, M. Watson, Bioinformatics 35, 2150 (2019).

111. P.-A. Chaumeil, A. J. Mussig, P. Hugenholtz, D. H. Parks, Bioinformatics (2019).

112. D. H. Parks, M. Imelfort, C. T. Skennerton, P. Hugenholtz, G. W. Tyson, Genome Research 25, 1043 (2015).

113. M. R. Olm, C. T. Brown, B. Brooks, J. F. Banfield, ISME Journal 11, 2864 (2017).

114. B. D. Ondov, et al., Genome Biol 17 (2016).

115. G. Marçais, et al., PLoS Computational Biology 14, e1005944 (2018).

116. M. D. Lee, Bioinformatics 35, 4162 (2019).

117. D. Hyatt, et al., BMC Bioinformatics 11 (2010).

118. S. R. Eddy, PLoS Computational Biology 7, e1002195 (2011).

119. R. C. Edgar, Nucleic Acids Research 32, 1792 (2004).

120. M. N. Price, P. S. Dehal, A. P. Arkin, PLoS ONE 5, e9490 (2010).

121. G. P. Wagner, K. Kin, V. J. Lynch, Theory Biosci. 131, 281 (2012).

122. M. R. Gradoville, B. C. Crump, R. M. Letelier, M. J. Church, A. E. White, Front. Microbiol. 8 (2017).

123. J. T. Evans, V. J. Denef, mSphere 5, e00971 (2020).

124. L. Breiman, Machine Learning 45, 5 (2001).

125. M. N. Wright, A. Ziegler, J. Stat. Soft. 77 (2017).

126. P. Probst, M. Wright, A.-L. Boulesteix, Wiley Interdisciplinary Reviews: Data Mining and Knowledge Discovery (2018).

127. D. J. Richter, C. Berney, J. F. H. Strassert, F. Burki, d. C. Vargas, bioRxiv p. 2020.06.30.180687 (2020).

128. S. Janitza, E. Celik, A.-L. Boulesteix, Adv Data Anal Classif 12, 885 (2016).

129. F. Degenhardt, S. Seifert, S. Szymczak, Briefings in Bioinformatics 20, 492 (2017).

130. Z. Šidák, Journal of the American Statistical Association 62, 626 (1967).

131. G. Csardi, T. Nepusz, InterJournal Complex Systems, 1695 (2006).

132. R. C. Team, R Foundation for Statistical Computing, Vienna, Austria: USBN pp. 3–900051 (2019).

133. S. Pesant, et al., Sci Data 2 (2015).

