## Supplementary Information for "Eukaryotic genomes from a global metagenomic dataset illuminate trophic modes and biogeography of ocean plankton"

Molecular and genomic approaches that target mixed microbial communities (e.g., metagenomics or metatranscriptomics) provide insight into the ecological roles, evolutionary histories, and physiological capabilities of the microorganisms and the processes in the environment. Computational tools that harness large-scale sequence surveys have become a valuable resource for characterizing the genetic make-up of the bacterial and archaeal component of the marine microbiome. Yet, fewer studies have focused on the unicellular eukaryotic fraction of the community. Here, we developed the EukHeist automated computational pipeline, to retrieve eukaryotic and prokaryotic metagenome assembled genomes (MAGs). We applied EukHeist to the eukaryote-dominated large-size fraction data (0.8-2000$\mu m$) from the *Tara* Oceans survey to recover both eukaryotic and prokaryotic MAGs, which we refer to as TOPAZ (*Tara* Oceans Particle-Associated MAGs). The TOPAZ MAGs consisted of more than 900 eukaryotic MAGs representing environmentally-relevant microbial and multicellular eukaryotes in addition to over 4,000 bacterial and archaeal MAGs. The bacterial and archaeal TOPAZ MAGs retrieved with EukHeist complement previous efforts by expanding the existing phylogenetic diversity through the increase in coverage of many likely particle- and host-associated taxa. We also demonstrate how the novel eukaryotic genomic content recovered from this study might be used to infer functional traits, such as trophic mode. By coupling MAGs and metatranscriptomic data, we explored ecologically-significant protistan groups, such as the Stramenopiles. A global survey of both eukaryotic and prokaryotic MAGs enabled the identification of ecological cohorts, driven by specific environmental factors, and putative host-microbe associations. Accessible and scalable computational tools, such as EukHeist, are likely to accelerate the identification of meaningful genetic signatures from large datasets, ultimately expanding the eukaryotic tree of life.

### 1 Comparison of ANI deduplication and MUMmer

#### 1.1 Explanation of use of fastANI

We used the fastANI tool via the CoverM software to deduplicate both eukaryotic and prokaryotic MAGs prior to abundance quantification in the raw reads, as detailed in the Materials and Methods. This implementation of fastANI uses the dRep formula for assessing genome quality (*1*); we used a percent identity threshold of 99% for dereplication and a 50% minimum coverage in order to reduce redundancy while retaining population-specific genes (*2*). In order to evaluate the impact of using different tools and clustering thresholds on the total number of eukaryotic MAGs reported by our study, we also applied the dnadiff tool from the MUMmer package (*3*) to the eukaryotic genomes Figure S25. While fastANI did tend to estimate a higher total number of genomes than dnadiff did at the same level of reference coverage and percentage identity Figure S25A, there was overall tight correspondence between the outcomes predicted by the two tools. Hence, we used fastANI as implemented in CoverM to dereplicate simultaneously with abundance quantification (https://github.com/wwood/CoverM). Similar fastANI parameters have also been used by previous MAG studies to differentiate at the strain level (e.g. (*4, 5*)).

### 2 Development and performance of the trophic mode model

#### 2.1 Description of the model and Heterotrophy Index

We used a variable selection algorithm and Random Forest machine learning model framework in order to predict the likely trophic mode of the eukaryotic TOPAZ MAGs described in this study. Transcriptomes from the MMETSP and EukProt were manually-annotated as phototroph, mixotroph, or heterotroph based on the literature (Supplementary Table 5). We tested our model with a randomly subset test set comprised of the 25% of MMETSP and EukProt transcriptomes (*6, 7*) that were excluded from the model building procedure. With this test subset we obtained an accuracy of 94.6% (Figure S16), meaning that nearly 95% of taxonomic annotations derived from the machine learning model aligned with their manually-assigned trophic mode annotation (Figure S16). When applied to the TOPAZ MAGs, all MAGs were either classified as phototrophs or heterotrophs, with none classified as mixotrophs. This likely reflects that the model was generally conservative when it came to assigning genomes or transcriptomes as mixotrophs (Figure S20). As a consequence, we developed a secondary metric for assessing the extent of heterotrophy in the test genomes and transcriptomes using the KOs selected by the vita selection process ($n = 1787$), but instead of using the presence or absence of these KOs as a binary indicator to inform the classification of the MAGs, we used the presence or absence as part of an equation to more sensitively assess the number of KOs present that tend to be indicative or either heterotrophy or phototrophy. The result was the "Heterotrophy Index" (H-index), a metric for assessing trophy based on KEGG pathway presence or absence.

The H-index is a sliding scale that weights the presence of heterotrophy, phototrophy- and mixotrophy-indicative KOs to assess the overall likely trophic state of an organism, and will consequently better show when an organism is more likely mixotrophic or possessing traits from both heterotrophy and phototrophy. Because mixotrophs were less common in our test dataset, there are natural concerns about the skill of the model when it comes to identifying them (*8*). We did not attempt to address the imbalance of the categorical training data in the model as other papers have recently explored (*9, 10*), hence our model retains the bias of reduced sensitivity when the distribution of the training data categories cannot be compared directly to the "true" incidence of mixotrophy among eukaryotic organisms (*11*). Because the majority of test and training transcriptomes were phototrophs or heterotrophs, it is more conservative for a Random Forest model to assign these modes more frequently. As the TOPAZ MAGs covered lineages with known mixotrophic members (*12*), and with comparison and feedback from an alternative trophic model as described in Section 2.2, we applied the H-index to provide more sensitivity in the identification of likely mixotrophs. In particular, with the Random Forest design, if a MAG has characteristics of both heterotrophs and phototrophs that are present in the training data, but does not align with the limited sample of mixotroph transcriptomes (which is also problematic due to the opportunistic nature of the sampling, see Section 2.3), these MAGs would be assigned the best guess between phototrophy and heterotrophy, when in reality this combination of traits may indicate some form of mixotrophy. The H-index also serves as a confidence metric for the trophy estimate. For example, a MAG with a large positive H-index would more confidently be called a heterotroph, as this would indicate a strong frequency and degree of alignment with heterotroph references (and, specifically, alignment with those heterotrophic references for KOs identified by the vita algorithm as important for distinguishing trophic mode within the training set). By contrast, a small (or near zero) positive H-index may be mixotrophic or represent a less complete MAG.

#### 2.2 Comparison to Burns and Lambert models

In order to assess the performance of our model, which relies solely on assessment of KOs (*13*) that were determined computationally to be important and assessed for function after the fact (Figure S18), we applied the model from (*14*) (heretofore referred to as the Burns model) to the same highly complete eukaryotic MAGs, as well as to the MMETSP transcriptomes. This model assigns a score from zero to one for individual characteristics related to trophy, including photosynthetic ability, phagocytosis, and prototrophy (*14*). Using Hidden Markov Models for selected genes which have known association with the aforementioned trophic strategies, the Burns model instead looks for a set of genes consistent with each trait, to assess the "completeness" of the genome or transcriptome with respect to the machinery known to be involved with each function. We found the Burns model results to be consistent with our H-index procedure in the following ways. Among the MMETSP transcriptomes, 93.1% (n=81) of the transcriptomes with a positive H-index (indicative of net heterotrophy) also had a photosynthesis score of less than 0.5 as assigned by the Burns model, and 90.8% (n=79) had a photosynthesis score of less than 0.05 via the Burns model (Figure S19). Similarly, 87.3% (n=226) of MMETSP transcriptomes with a negative H-index (indicative of net phototrophy) had a photosynthesis score of greater than 0.5 as assigned by the Burns model (52.9% (n=137) had negative H-index and Burns photosynthesis score greater than 0.95). MMETSP transcriptomes with a zero or near-zero H-index, which corresponds to putative mixotrophy, had varying photosynthesis scores according to the Burns model, but were more likely to have high (0.6-1) photosynthesis scores, which is consistent with mixotrophy (Figure S19). However, several of the MAGs which were predicted to be mixotrophs by the Random Forest model and were annotated manually as mixotrophs from the available metadata had mid- to high- photosynthesis scores in the Burns model, yet negative-leaning H-index scores (Supplementary Table 10). These transcriptomes also tended to have high (>0.7) phagocytosis predictions per the Burns model, consistent with the presence of genetic resources for both heterotrophic and phototrophic strategies. Broadly, we found that, similarly to our H-index and

Random Forest model annotations, the Burns photosynthesis prediction results tended to align with the expected lifestyle of each MAG based on EUKulele (*15*) taxonomic annotations, with expected heterotrophs like Amoebozoa, Fungi, and Opisthokonta scoring low on photosynthetic ability, while expected phototrophs like Ochrophyta and Chlorophyta tended to score highly for photosynthetic ability (Figure S19). Most disagreement was found within the SAR clade and Crypophyta, wherein a range of photosynthesis scores were found by the Burns model, and sometimes these scores contradicted the annotation found by the H-index and Random Forest model (fig. S19 and Supplementary Tables 9 and 10). This would indicate potentially cryptic and variable trophic strategies and lifestyles within these annotated groups.

When split into classes of photosynthetic ability based on Burns model scores (to isolate "highly" or "not at all" photosynthetic: 0-0.1, 0.1-0.45, 0.45-0.55, 0.55-0.9, 0.9-1), MMETSP transcriptomes with "no" photosynthesis according to the Burns model (photosynthesis prediction $< 0.1$) had an average H-index of $38.85 \pm 68.19$, while transcriptomes with "high" photosynthesis (photosynthesis prediction $> 0.9$) had an average H-index of $-272.34 \pm 119.94$ (Figure S17). As far as the TOPAZ MAGs, we similarly found that the MAGs predicted to be heterotrophic had low variance in photosynthesis prediction as reported by the Burns model, yet the variability in the photosynthetic prediction was high, in particular among those MAGs of higher (less negative, hence closer to "mixotrophic") H-index (Figure S20).

#### 2.3 The future of trophic mode models

The model we developed relies solely upon references that were derived from expression-level data (transcriptomics). Additionally, we used the entire MMETSP (*6*) and EukProt (*7*) databases with manually assigned trophic strategies based on the literature (Supplementary Table 5). Both of these choices carry issues that may be responsible for our under prediction of mixotrophy. First, as they were transcriptomic datasets, the experimental conditions that were used to generate the reference transcriptome are important, in that if a culture was maintained in phototrophy-favorable conditions (e.g. high light, sufficient inorganic nutrients) as opposed to heterotrophy-favorable conditions (e.g. low light, external carbon source), the transcripts reconstructed from these experiments may result in a reference that is skewed towards phototrophy or heterotrophy, respectively. Regardless of the conditions in which the organism was grown, it is likely that genes present in the genome of the organism were not recovered by transcriptomics. While this means that important genes related to trophic strategy may be missed from the prediction workflow, this is also exciting, as there is much room for growth for these already accurate and skillful models. As genomic references become available and the number of transcriptomic experiments continue to grow, we can expect to further constrain the classes of genes that decide trophic mode. As databases grow, they may be pruned to include only experiments in which the trophy of the organism was known exactly, which would enable the patterns of expression which decide trophic mode to be pinpointed more precisely. Ideally, a complete core set of protein families necessary for heterotrophy and for phototrophs could be identified. A fundamental question that remains, however, is if there are any genes or protein families that are characteristic of mixotrophic organisms *only* or if these organisms are better characterized based on the co-occurrence of genes indicative of phototrophy and heterotrophy. If the latter, mixotrophy would be best characterized by the proportion of these sets which overlap in the genome or the expression profile of the candidate organism. While this is in principle the aim of the Burns model ( (*14*), Section 2.2), this approach might still be augmented by machine learning-based techniques that have the capacity to identify important genes that may not have yet been annotated.

### 3 Correlations with environmental parameters

The relative abundance of eukaryotic MAGs across samples was correlated with environmental parameters (Figure S12) and were found to broadly cluster by course taxonomic grouping, as such abundances were summed by broad taxonomic group to assess (Figure S13) general trends and correlations in patterns of abundance. Globally, we note that metazoan MAG abundance significantly (Bonferroni corrected $p < 0.001$) positively correlated with temperature, salinity, and retention time (defined as the average length of time that a particle has been trapped in an eddy), and significantly negatively correlated with a variety of nutrients (phosphate, nitrate+nitrite, silica), CDOM, and depth (meaning they are more abundant in surface samples). This suggests that the metazoan MAGs we are recovering are likely abundant in the subtropical and tropical waters of the large ocean gyres that are oligotrophic. Notably, Choanozoa and Amoebozoa clustered with Metazoans, forming a distinct cluster from other more typically photosynthetic and planktonic groups. Chlorophyta, Apusozoa, Cryptophyta, Dinoflagellata, Ochrophyta, SAR, and Haptophyta clustered separately from Metazoa and were typified by significant positive correlation with oxygen, $a$, and nitrite (Figure S13). Additionally, MAGs recovered that belonged to Ochrophyta were found to be significantly negatively correlated with temperature. This suggests that these MAGs were likely associated with more polar or subpolar regions.

### Supplementary Tables

All Supplementary tables are available through the open science framework at `https://osf.io/twz2f/`.

- **Supplementary Table 1.** Assembly group description, sample inclusion, and basic assembly statistics.
- **Supplementary Table 2.** TOPAZ Eukaryotic MAG taxonomy, genomic characteristics (e.g., total length, GC content, N50, number of predicted proteins,), and estimated completeness and contamination.
- **Supplementary Table 3.** TOPAZ Prokaryotic MAG taxonomy as estimated by GTDB, dataset indication (non-redundant representatives (NR), total size, and estimated completeness and contamination.
- **Supplementary Table 4.** TOPAZ Prokaryotic MAG summary of recovered MAGs across phyla. Total counts are shown for all MAGs (All), the non-redundant subset of MAGs (AllNR), and the high-quality, non-redundant MAGs (HQNR).
- **Supplementary Table 5.** Reference transcriptomes used in the construction and testing of the trophic model. The manually annotated trophic status and details, relevant reference literature, and taxonomic information are noted. Additionally, a presence/absence matrix is provided for all eukaryotic KEGG ids considered here.

- **Supplementary Table 6.** Vita selected KOs and their associated heterotrophy, phototrophy, and mixotrophy ratios as described in equations 1-4.
- **Supplementary Table 7.** KEGG presence and absence for the TOPAZ eukaryotic MAGs, as was used in the trophic model.
- **Supplementary Table 8.** Pfam presence and absence across the TOPAZ eukaryotic MAGs.
- **Supplementary Table 9.** Eukaryotic TOPAZ MAG predicted trophic status and heterotrophy index (H-index).
- **Supplementary Table 10.** Burns model (*14*) predicted prototrophy, photosynthetic ability, and phagocytosis for the TOPAZ eukaryotic MAGs.
- **Supplementary Table 11.** Network analysis community composition.
- **Supplementary Table 12.** Eukaryotic cluster groups derived from average nucleotide identity clustering of eukaryotic TOPAZ MAGs with the Delmont eukaryotic MAGs (*16*) based on an ANI cutoff of 99%.
- **Supplementary Table 13.** Environmental correlations with network derived communities.

### Supplementary Figures

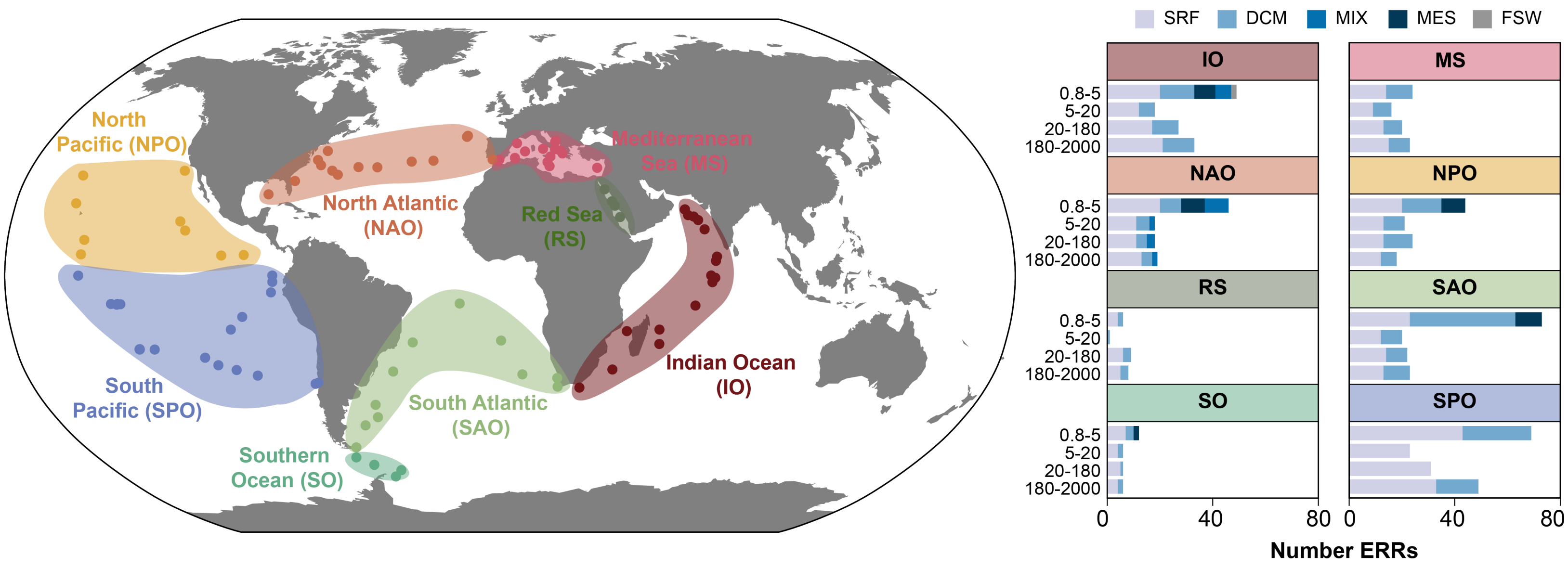


**Figure S1. Sample map and distribution of sequence data sets across regions.** Stations from *Tara* Oceans were categorized into general regions which are highlighted by color. The depth (surface (SRF), deep chlorophyll max (DCM), mixed surface sample (MIX), mesopelagic (MES), and filtered seawater (FSW)) and size fraction are shown as bar plots for each ocean region.

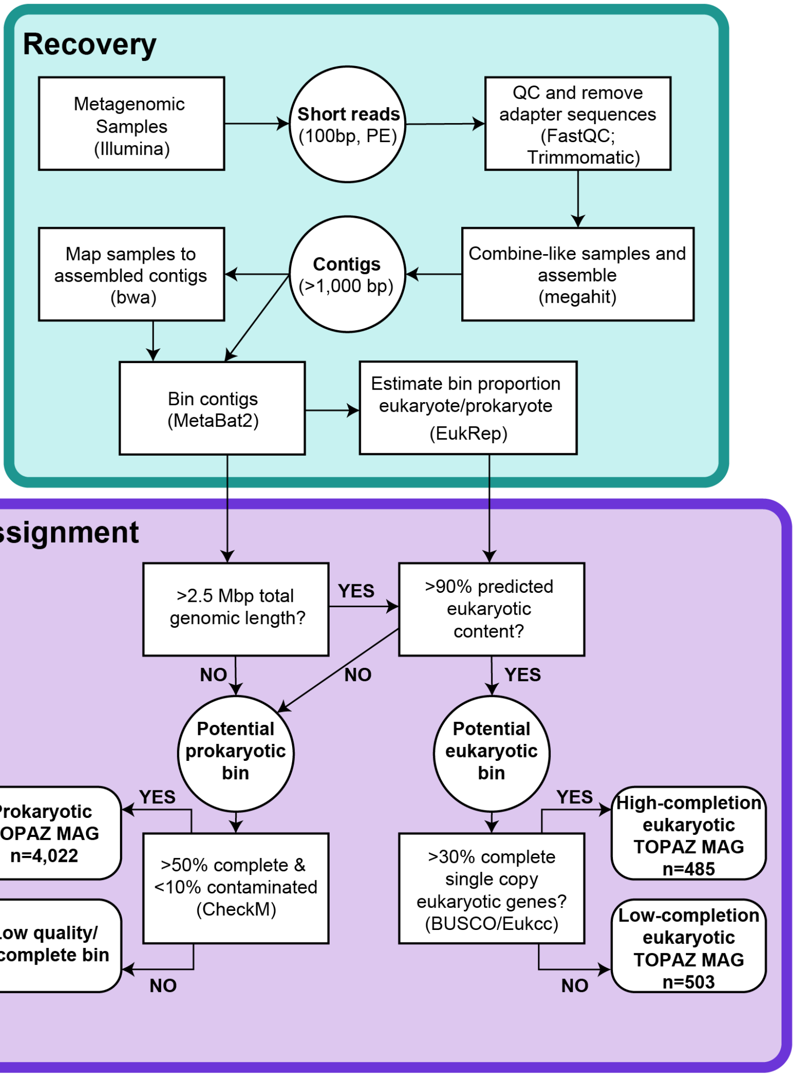


**Figure S2. Flow chart of the EukHeist pipeline.** Workflow for EukHeist supported through Snakemake (https://github.com/AlexanderLabWHOI/EukHeist).

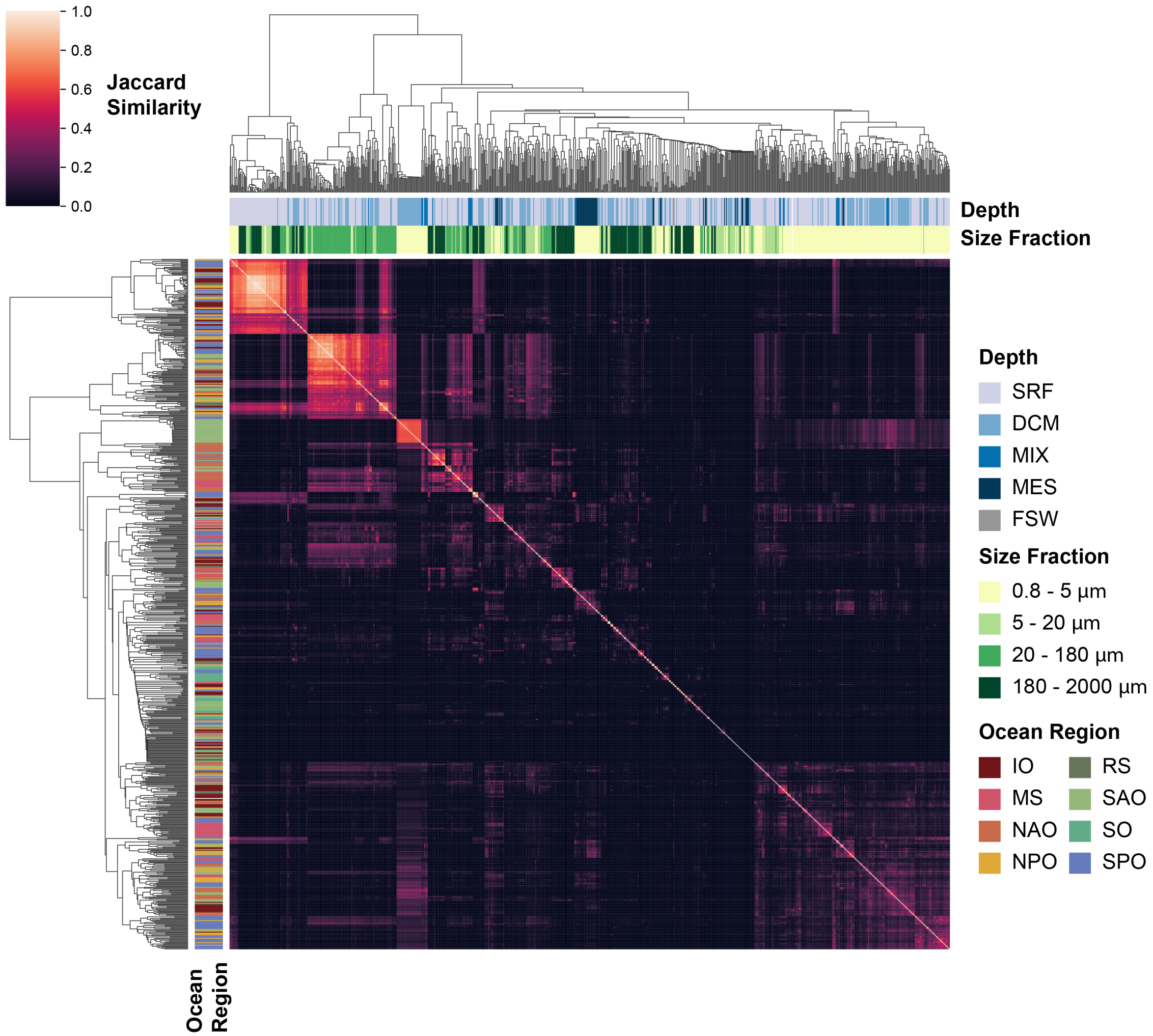


**Figure S3. Sourmash comparison of all metagenomic samples from *Tara* Oceans large size fraction dataset.** A minhash comparison was calculated using sourmash (k=31, scale=10,000) of the 824 metagenomic samples corresponding to the large size fraction metagenomic data from *Tara* Oceans (PRJEB4352) (*17*). The relative sequence content similarity is shown as Jaccard similarity. Hierarchical clustering of samples based on sequence content is shown and sample identity (sample depth, size fraction, and ocean region) is highlighted by colored blocks.

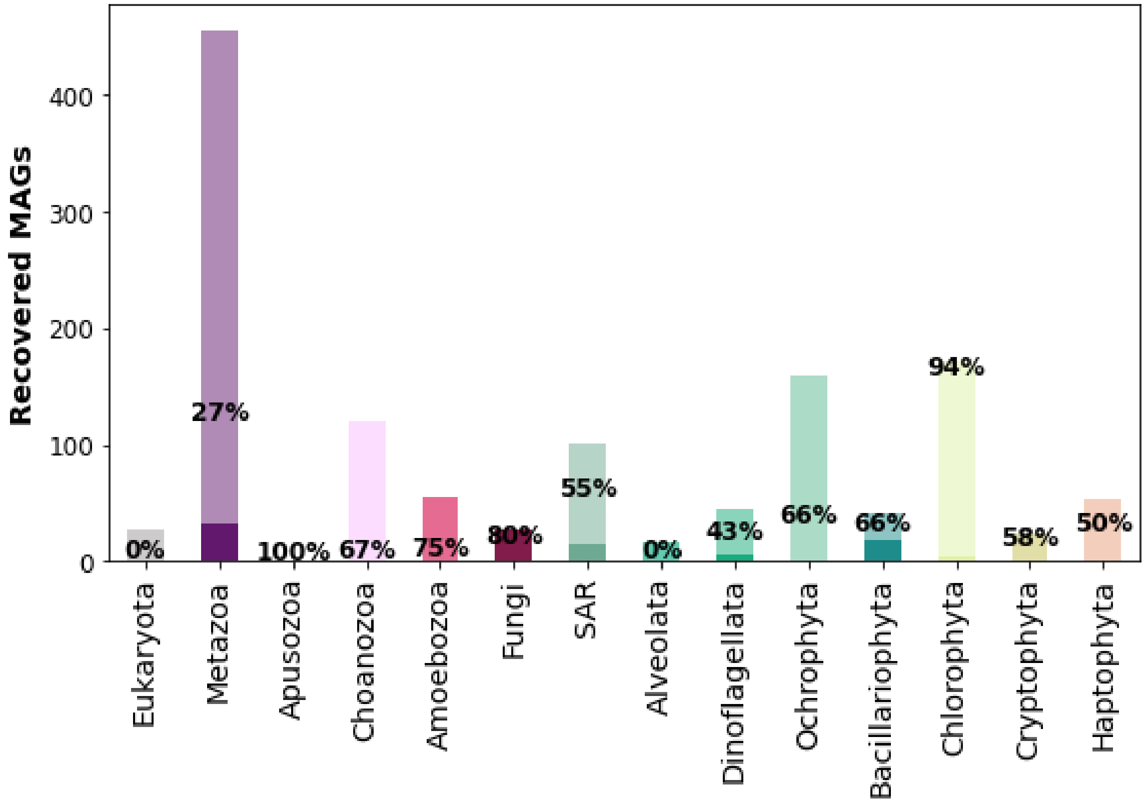


**Figure S4. TOPAZ Recovered Eukaryotic MAGs.** Course level taxonomic categorization of recovered eukaryotic TOPAZ MAGs (n=988). For each taxonomic group, the total number of MAGs is depicted. MAGs within a taxonomic group that were highly complete (> 30% BUSCO completeness) are shaded and the percentage of highly complete MAGs for each taxonomic group is reported.

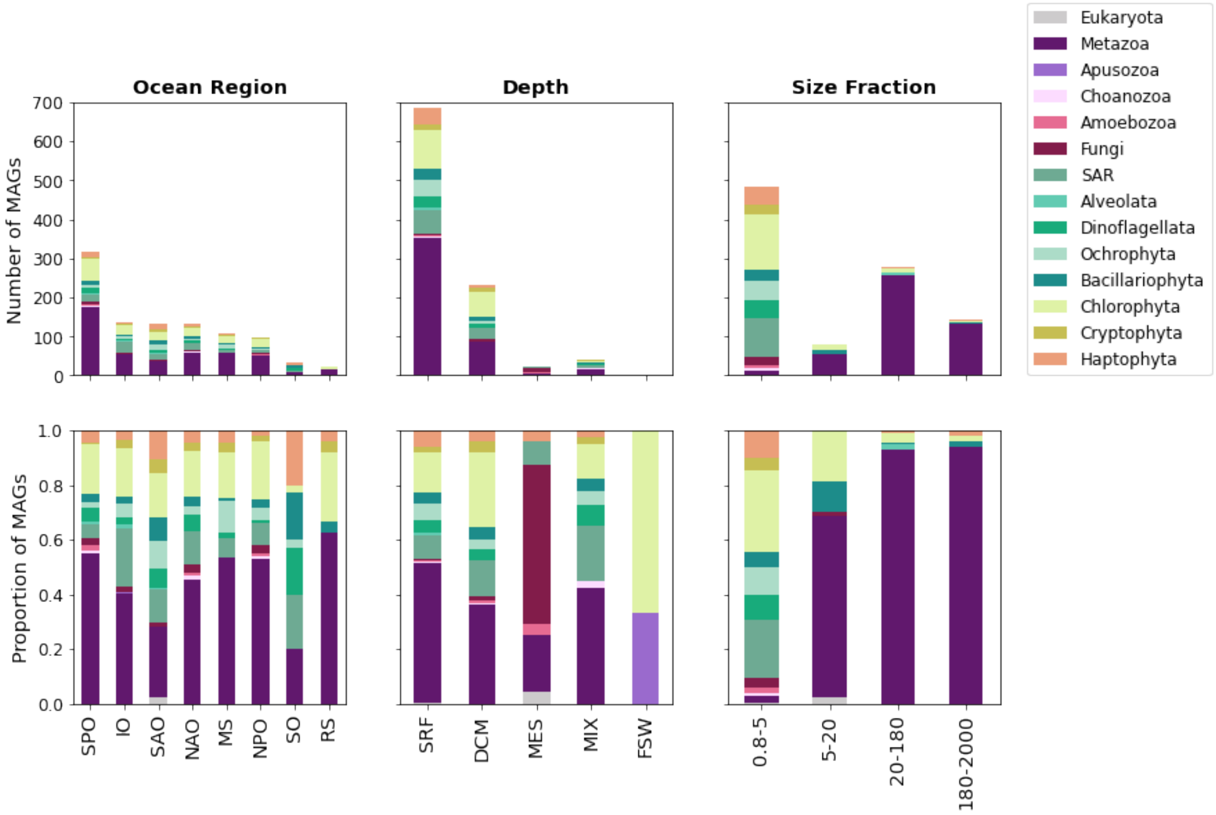


**Figure S5. TOPAZ Eukaryotic MAG as recovered by assembly group.** The taxonomic breakdown of eukaryotic MAGs recovered within each general type of assembly group (based on Ocean Region, Depth, and Size Fraction) for all eukaryotic MAGs recovered in this study (n=988). Taxonomy is shown both as a total number recovered (top) and as a proportion of MAGs recovered for a given category (bottom). Abbreviated ocean regions (*x*-axis of leftmost panels) follow the convention of the Tara Oceans dataset and are (left to right): South Pacific Ocean (SPO), Indian Ocean (IO), South Atlantic Ocean (SAO), North Atlantic Ocean (NAO), Mediterranean Sea (MS), North Pacific Ocean (NPO), Southern Ocean (SO), Red Sea (RS). Abbreviated general water column depths also follow the Tara Oceans convention and are (left to right): surface (SRF), deep chlorophyll maximum (DCM), mesopelagic zone (MES), marine epipelagic mixed layer (MIX), filtered seawater (FSW). Size fractions are to be interpreted as named, corresponding to size in micrometers. Taxonomic groups follow nomenclature elsewhere, with SAR comprised of Stramenopiles, Alveolates, and Rhizarians not covered under other taxonomic headings.

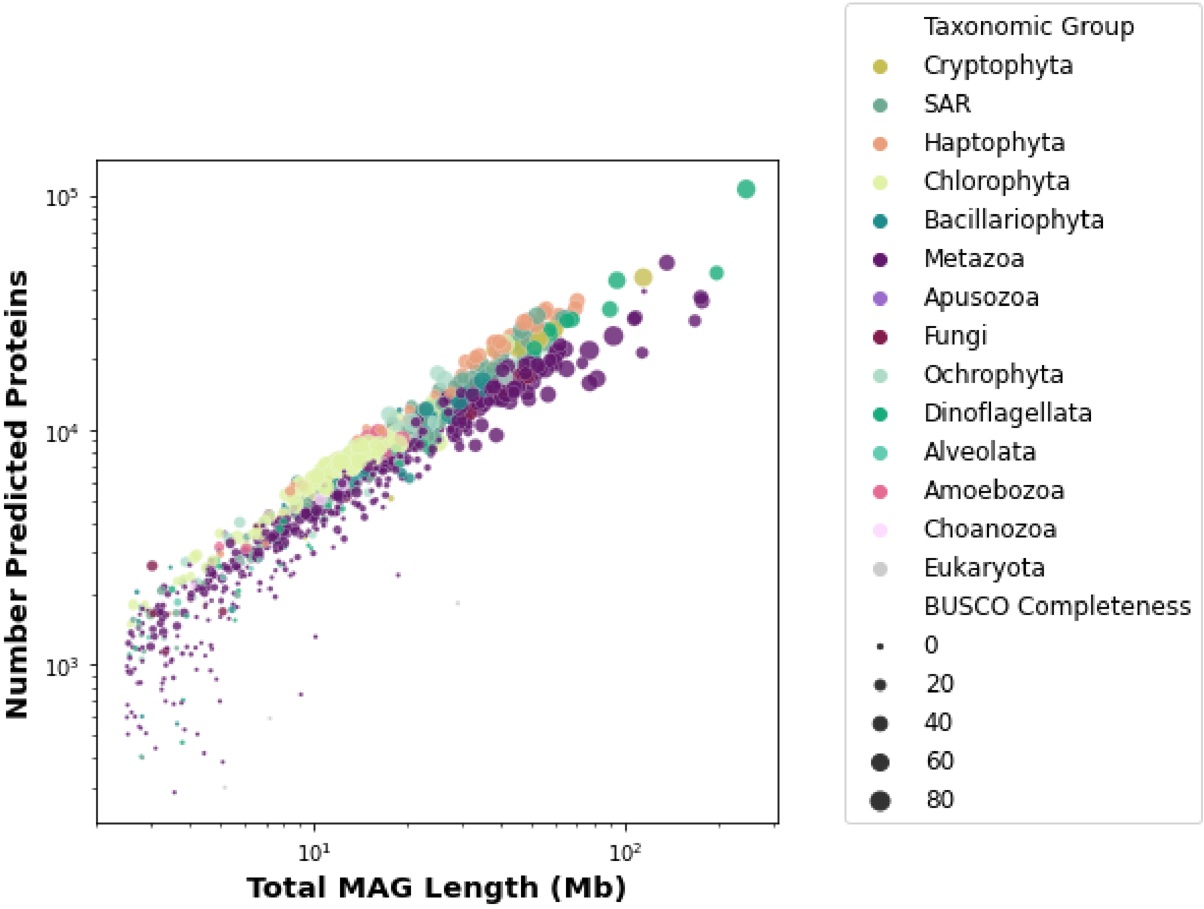


**Figure S6. The number of predicted proteins as a function of total MAG length.** The number of predicted proteins for each eukaryotic TOPAZ mag (n=988) is plotted against the total MAG length (Mb). Each MAG is colored by its taxonomic group and the size of the circle is scaled by the estimated BUSCO completeness.

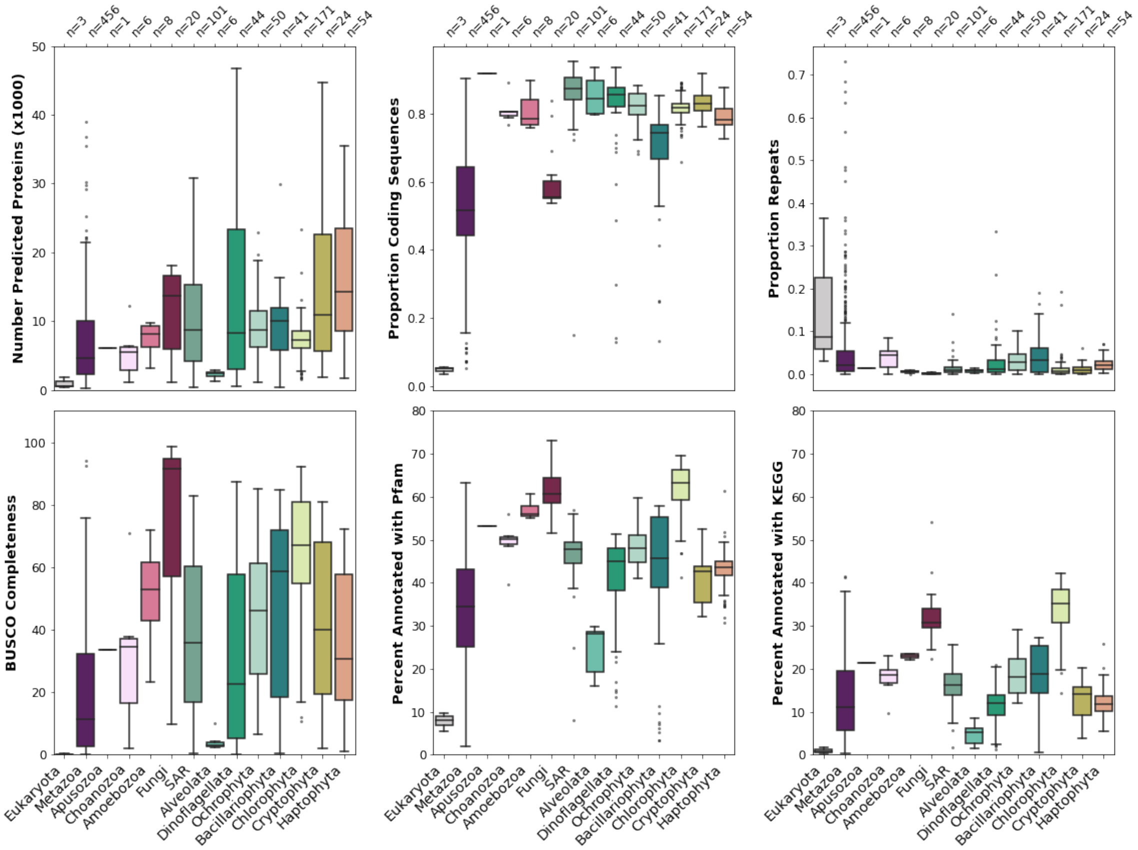


**Figure S7. Genomic traits of recovered eukaryotic completeness, protein predictions, and annotation of all eukaryotic TOPAZ MAGs (n=988).** The number of predicted proteins, proportion coding sequences, proportion repeat content, BUSCO completeness, percent annotation with Pfam and KEGG ontology are shown as box and whisker plots for the major higher-level groups that we define for this paper.

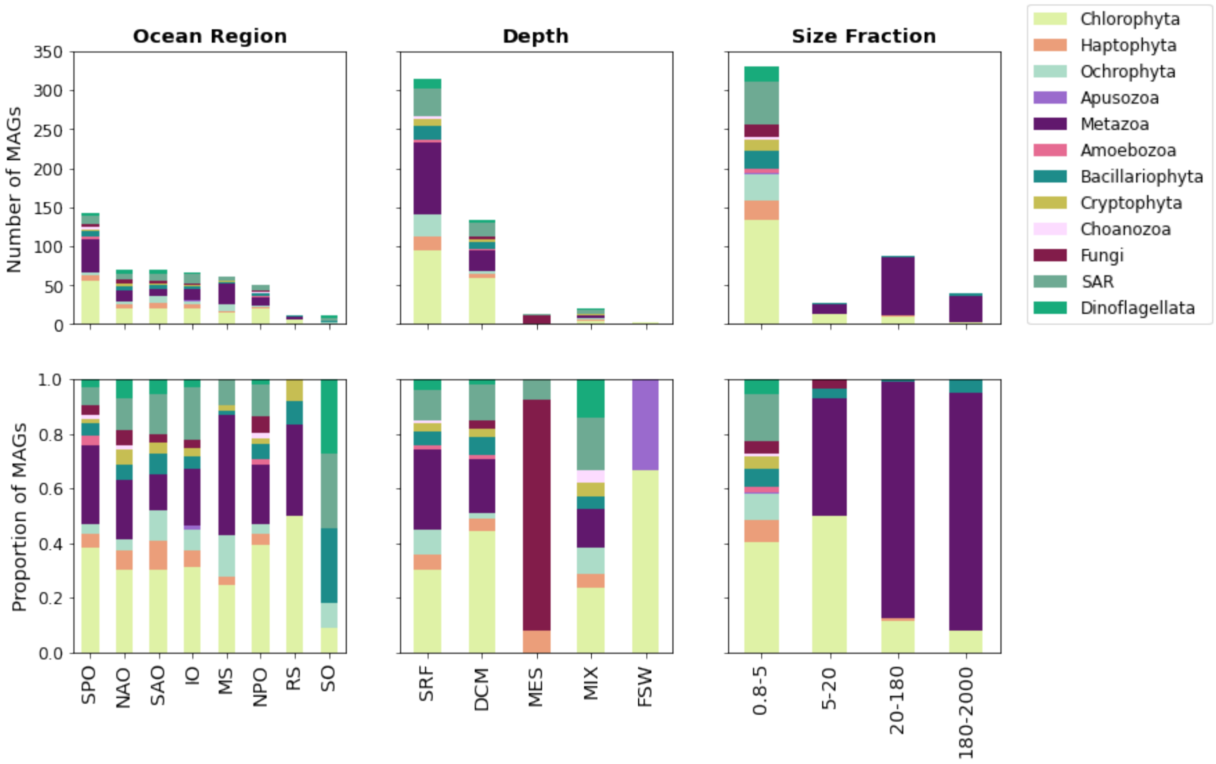


**Figure S8. TOPAZ Highly Complete Eukaryotic MAGs as recovered by assembly group.** The taxonomic breakdown of eukaryotic MAGs recovered within each general type of assembly group (based on Ocean Region, Depth, and Size Fraction) for highly complete eukaryotic MAGs (> 30% BUSCO completeness)recovered in this study (n=485)). Taxonomy is shown both as a total number recovered (top) and as a proportion of MAGs recovered for a given category (bottom).

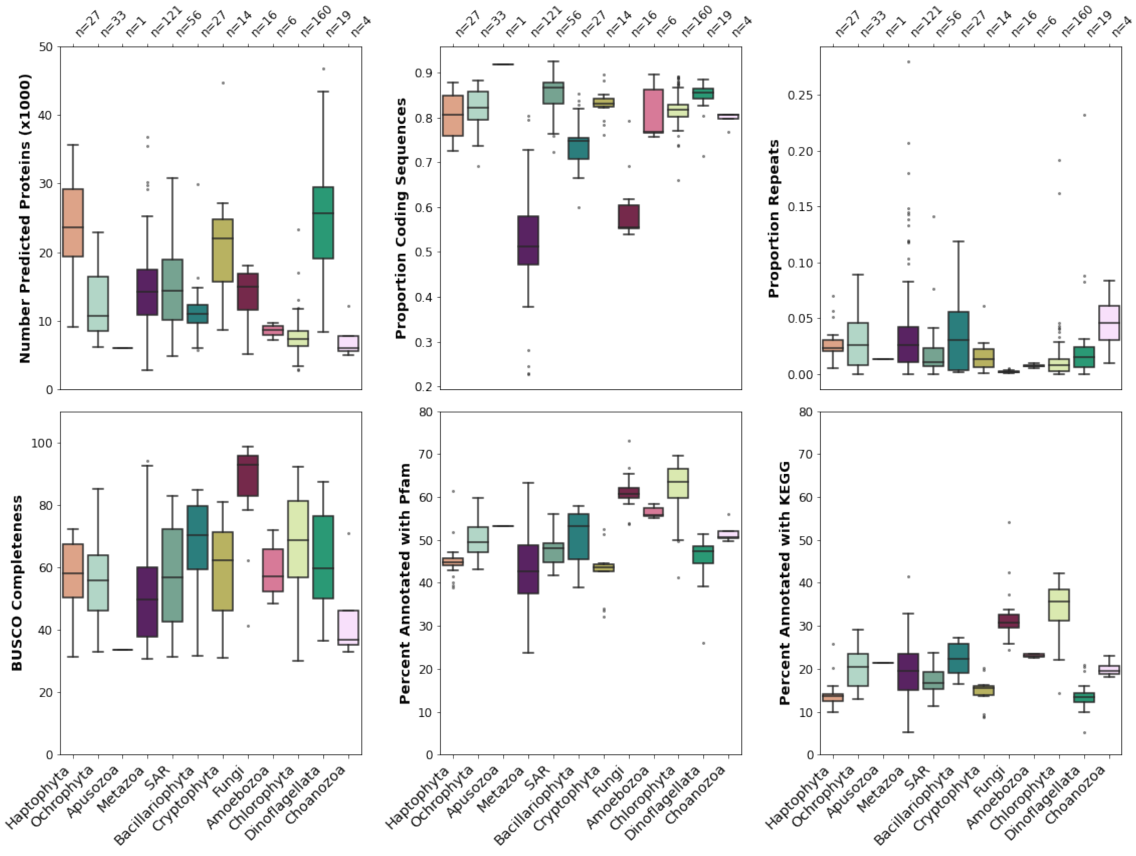


**Figure S9. Genomic traits of recovered eukaryotic completeness, protein predictions, and annotation of the highly complete eukaryotic TOPAZ MAGs (n=485).** The number of predicted proteins, proportion coding sequences, proportion repeat content, BUSCO completeness, percent annotation with Pfam and KEGG ontology are shown as box and whisker plots for the major higher-level groups that we define for this paper.

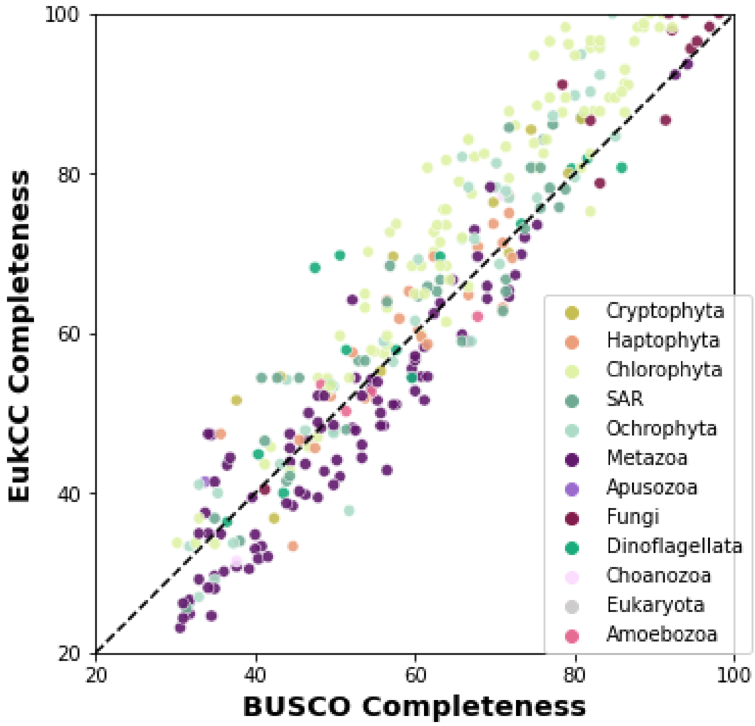


**Figure S10. A comparison of two metrics of eukaryotic completeness.** BUSCO completeness as defined based on the presence and absence of genes within the eukaryota_odb10 dataset was compared against the estimated EukCC completeness based on estimated lineages of given MAGs. Generally, it was observed that EukCC performed particularly well for certain groups (e.g. chlorophytes and fungi) and less well for others (e.g. metazoa).

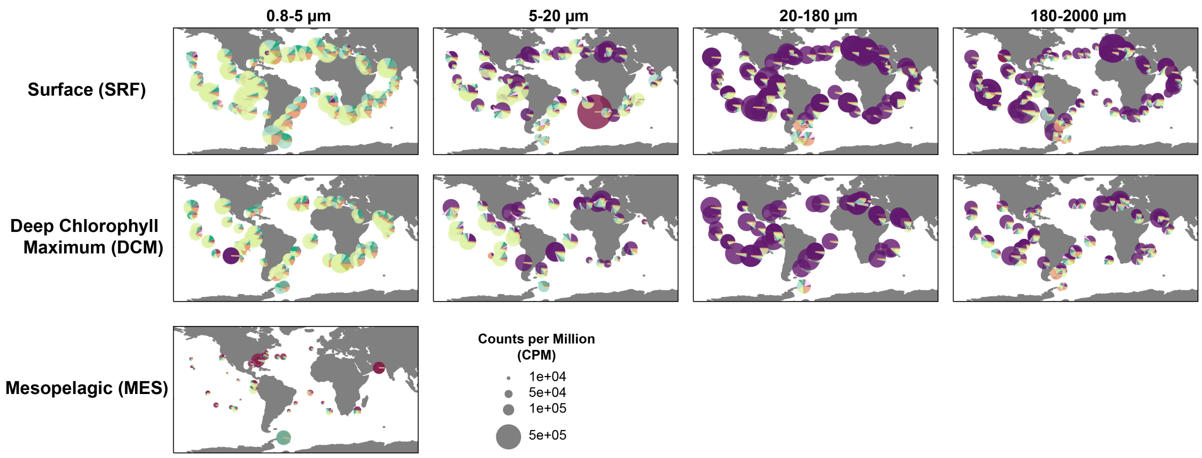


**Figure S11. The distribution of the major lineages of eukaryotic TOPAZ MAGs recovered across the *Tara* Oceans metagenomic datasets.** The counts per million (CPM) is depicted for all stations depicted by depth and size fraction.

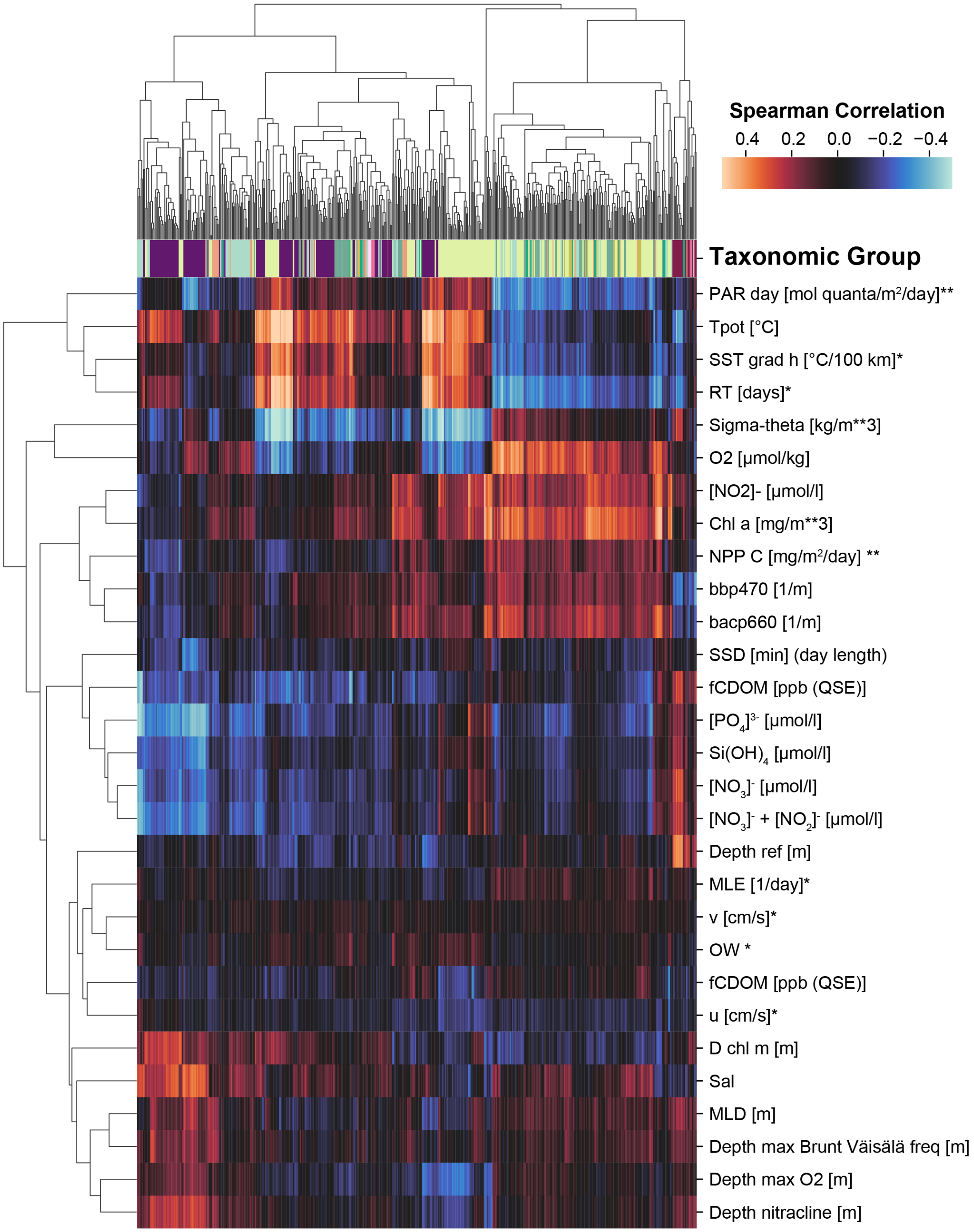


**Figure S12. A Spearman correlation between the metagenomic abundance of each of the 485 high-completion eukayrotic TOPAZ MAGs and tracked environmental parameters.** Environmental parameters include those directly from the sampling (*18*), modeled mesoscale physical features based on d'Ovidio et al. (*19*) (indicated with *), and averaged remote sensing products (indicated with **).

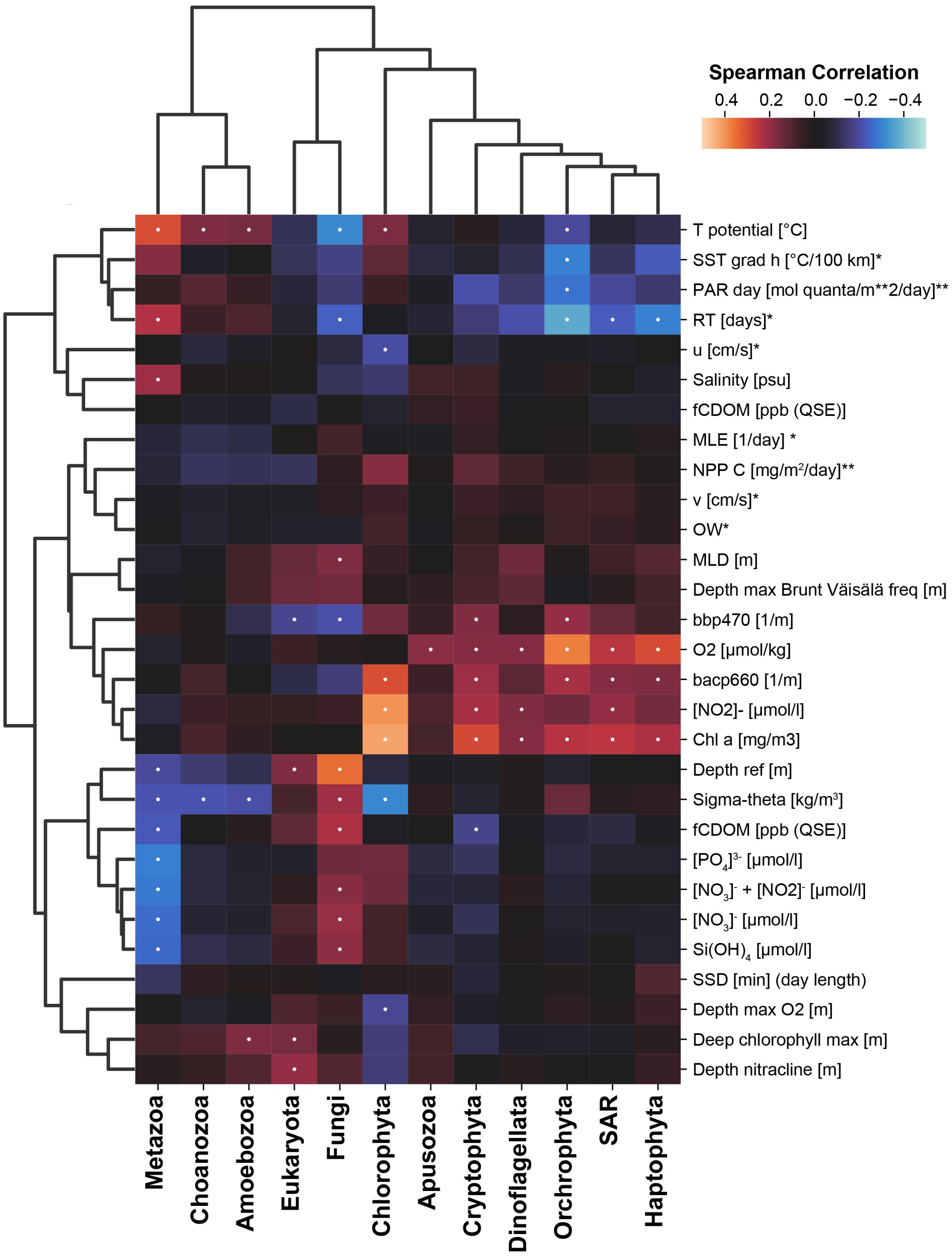


**Figure S13. A Spearman correlation between the summed metagenomic abundance of each taxonomic group and environmental parameters.** Environmnetal parameters include those directly from the sampling (*18*), modeled mesoscale physical features based on d'Ovidio et al. (*19*) (indicated with *) , and averaged remote sensing products (indicated with **). Significant Spearman correlations, those with a Bonferroni adjusted $p < 0.01$, are indicated with a dot on the heatmap.

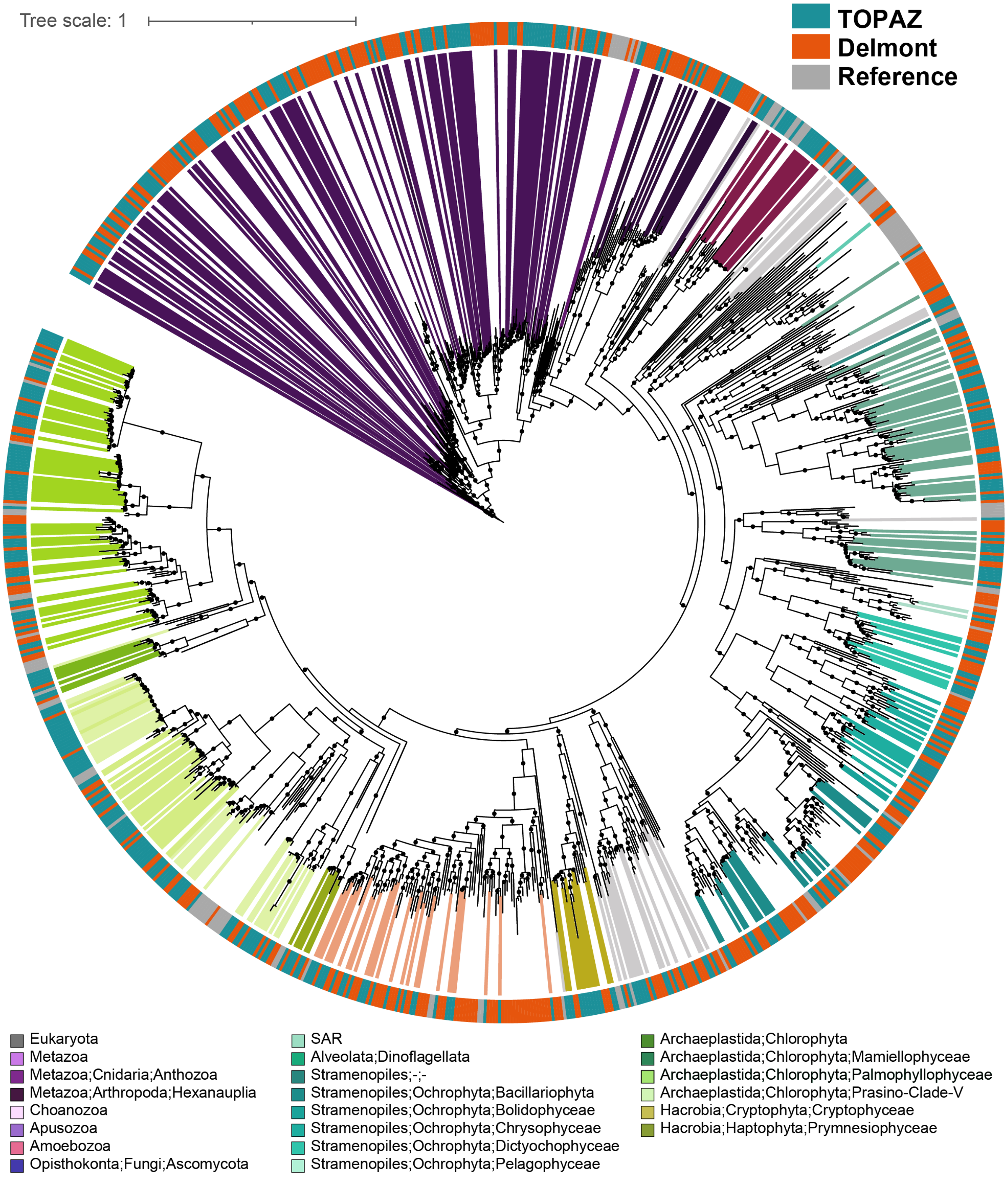


**Figure S14. A concatenated protein phylogeny containing TOPAZ and Delmont (*16*) eukaryotic MAGs.** MAGs that were estimated to be greater than 30% complete as well as reference genomes from cultured isolates. The maximum likelihood tree was inferred from a concatenated protein alignment of 49 proteins from the eukaryotic BUSCO gene set that were found to be commonly present across at least 75% of the 485 TOPAZ eukaryotic MAGs that were estimated to be >30% complete based on BUSCO ortholog presence (the same proteins that were used in Figure 1). Branches (nodes) corresponding to TOPAZ MAGs are colored based on consensus protein annotation estimated by EUKulele and MMSeqs.

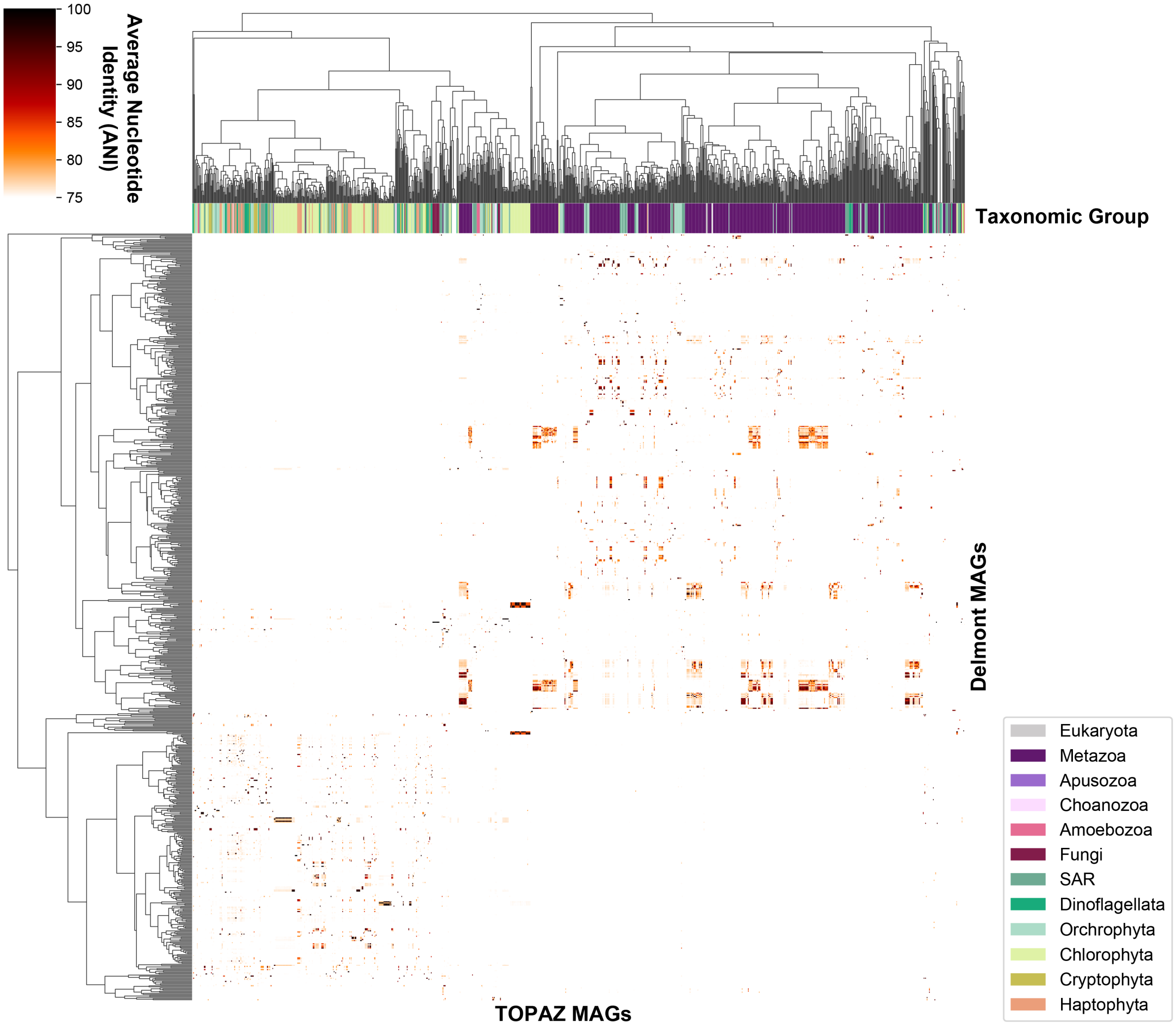


**Figure S15. Average nucleotide identity (ANI) between the High Completion TOPAZ and Delmont MAGs.** The ANI as estimated by FastANI between all eukaryotic TOPAZ MAGs and Delmont Eukaryotic MAGs is depicted. The taxonomic affiliation of the TOPAZ MAGs is depicted by color along the x-axis. A clustering was done with a cutoff of 99% ANI generating 83 unique clusters of MAGs, of which 46 contained a representative from both TOPAZ and Delmont (Supplementary Table 12).

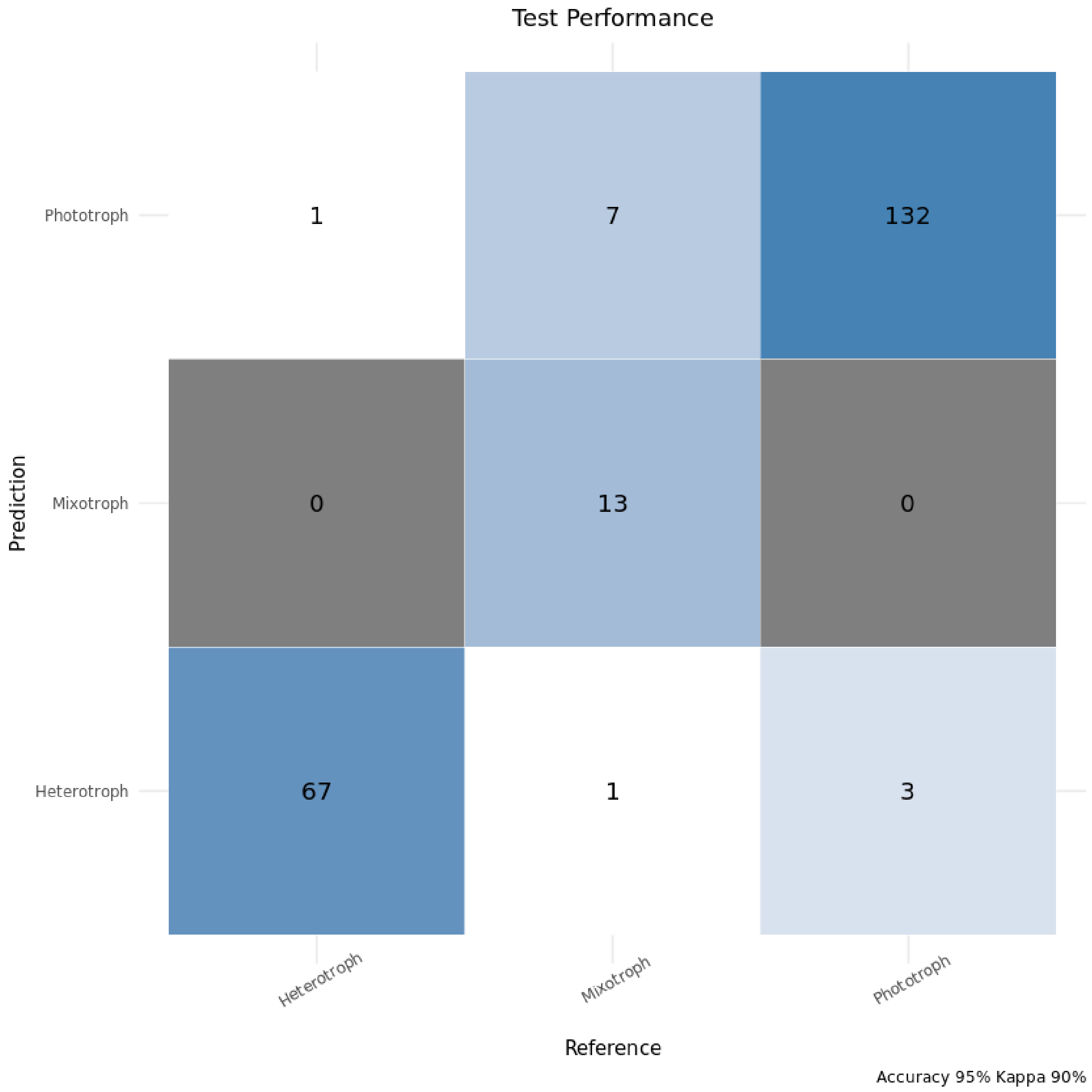


**Figure S16. Confusion matrix generated from Random Forest model construction on the 25% of the reference transcriptomes used for testing.** The *x* categorical axis is the manually-annotated trophic mode, while the *y* categorical axis is the predicted trophic mode via the Random Forest model. The greatest number of erroneous predictions involved mixotrophy (8 of 12).

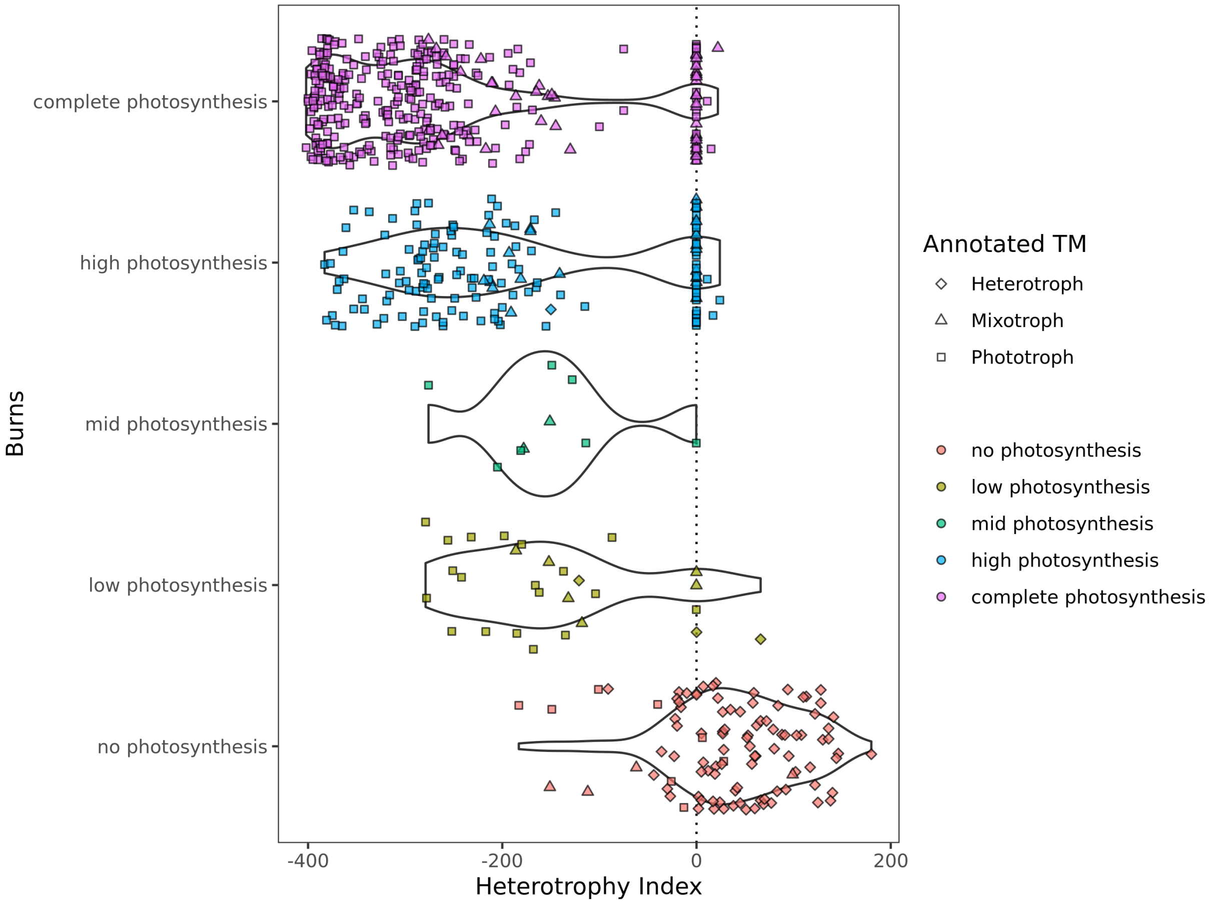


**Figure S17. Comparison of Heterotrophy Index scores with the Burns model prediction for photosynthetic ability.** On the *y* axis, MAGs are split into categories based on their photosynthesis score from the Burns model (which ranges from zero to one): "complete photosynthesis" (0.9-1), "high photosynthesis" (0.55-0.9), "mid photosynthesis" (0.45-0.55), "low photosynthesis" (0.1-0.45) and "no photosynthesis" (0-0.1).

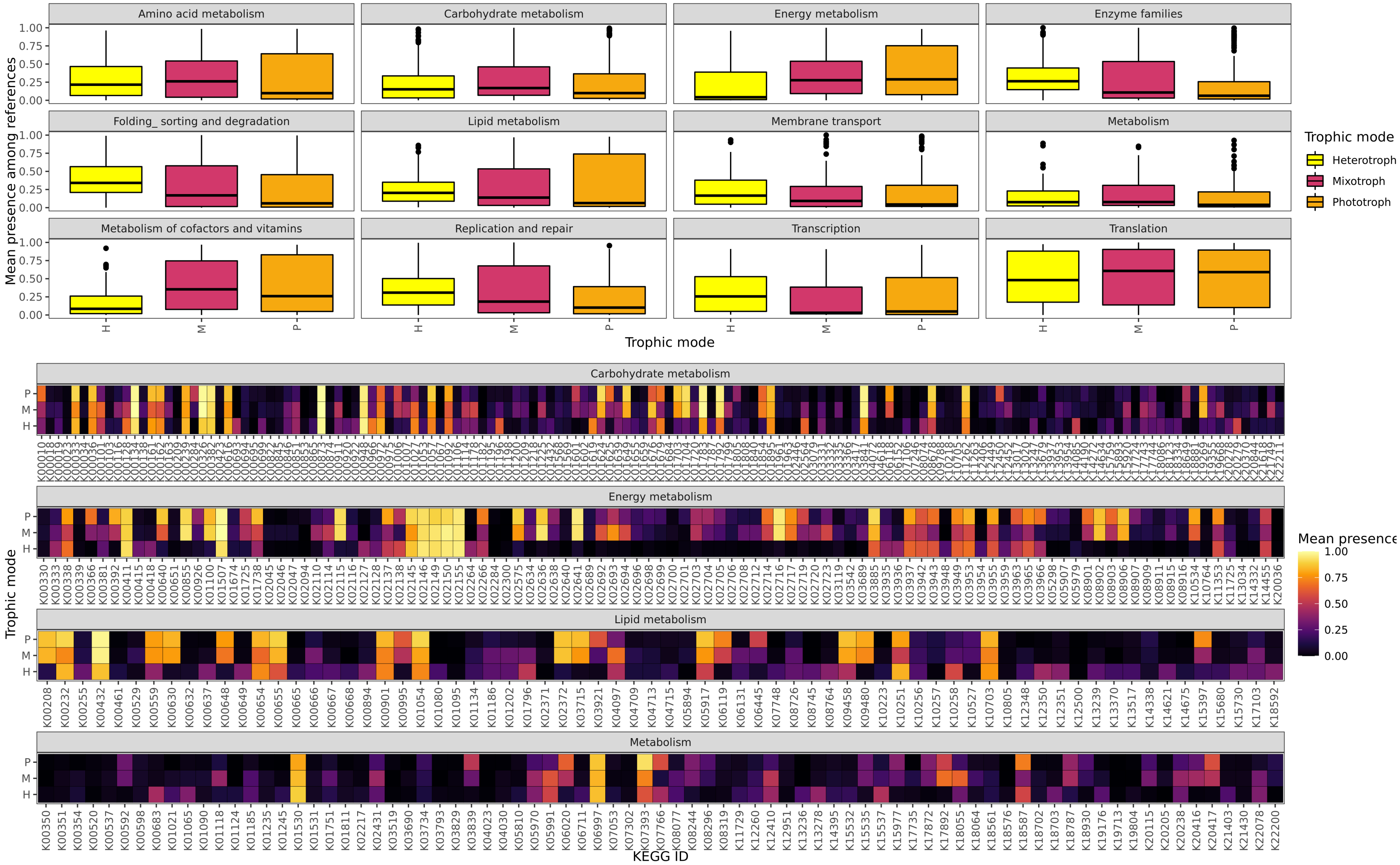


**Figure S18. Presence-absence of KEGG IDs as aggregated across all reference transcriptomes (MMETSP and EukProt) used in the trophic mode modeling process.** The top boxplot panel shows the mean presence of the KEGG IDs implicated in the listed pathways for each of the three identified trophic modes, while the bottom heatmap panel shows the average presence of each individual KO along the horizontal axis for reference transcriptomes annotated as each major trophic mode (vertical axis).

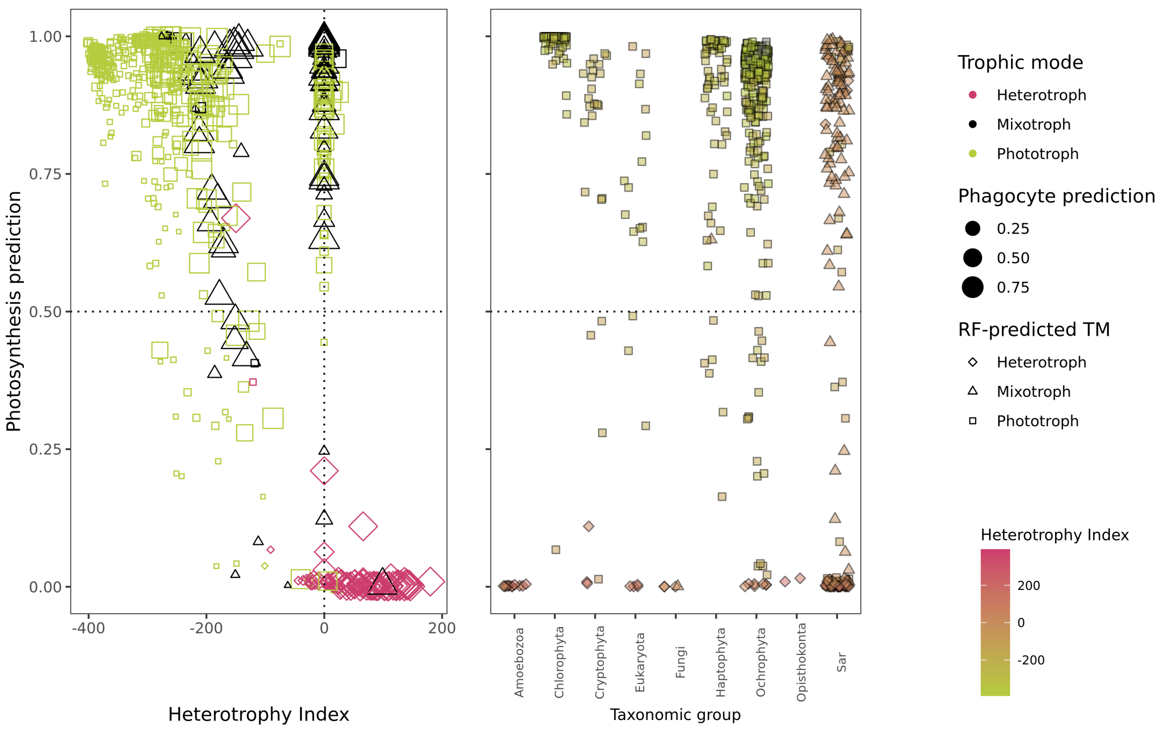


**Figure S19. Comparison of the Burns (*14*) model to our Random Forest predictions and Heterotrophy Index calculations for the reference MMETSP transcriptomes.** Left: Burns (*14*) photosynthesis predictions vs. composite Heterotrophy Index, scaled by the phagocytosis prediction and colored by the manually-annotated trophic mode. Right: predicted photosynthetic ability by taxonomic grouping, colored by the calculated Heterotrophy Index.

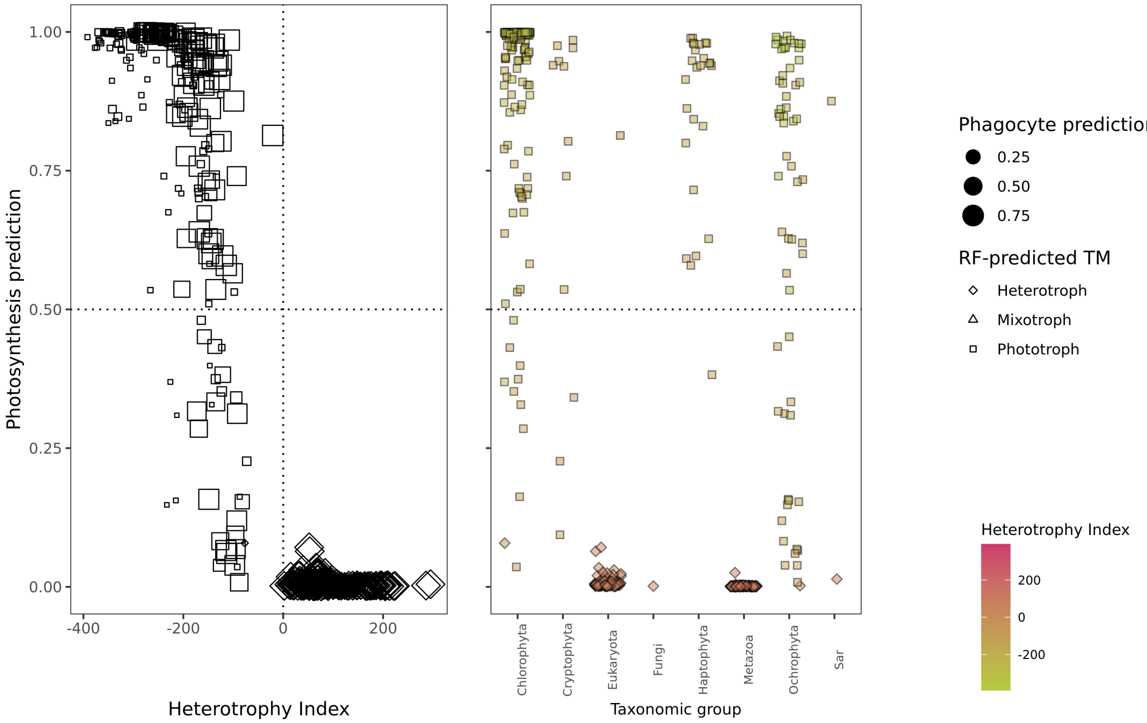


**Figure S20. Comparison of the Burns (*14*) model to our Random Forest predictions and Heterotrophy Index calculations for the TOPAZ MAGs.** Left: Burns (*14*) photosynthesis predictions vs. composite Heterotrophy Index, scaled by the phagocytosis prediction and shape indicating the Random Forest-derived predicted trophic mode (note there is no color because trophic mode could not be manually annotated). Right: predicted photosynthetic ability by taxonomic grouping, colored by the calculated Heterotrophy Index.

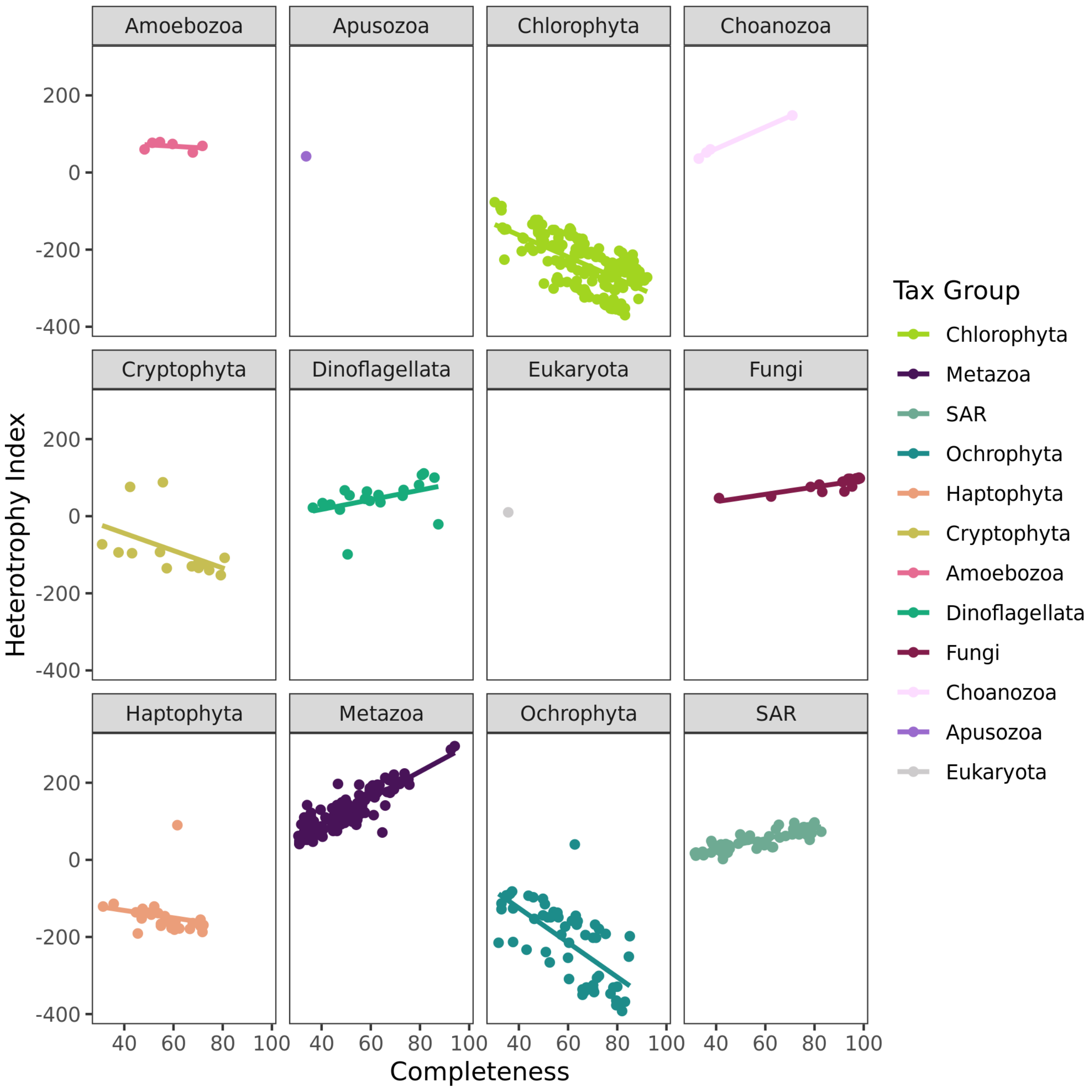


**Figure S21. Heterotrophy Index scores as a function of BUSCO completeness, faceted by taxonomy phylum.** The strongest association of Heterotrophy Index with completeness was found within Metazoa, for which it appears that the magnitude of the Heterotrophy Index is tightly coupled to the level of completeness of the MAG. By contrast, Cryptophyta (which are known to be mixotrophic (*12*)) showed a much weaker coupling of completeness with the magnitude of the Heterotrophy Index, and all values of the Index were closer to zero.

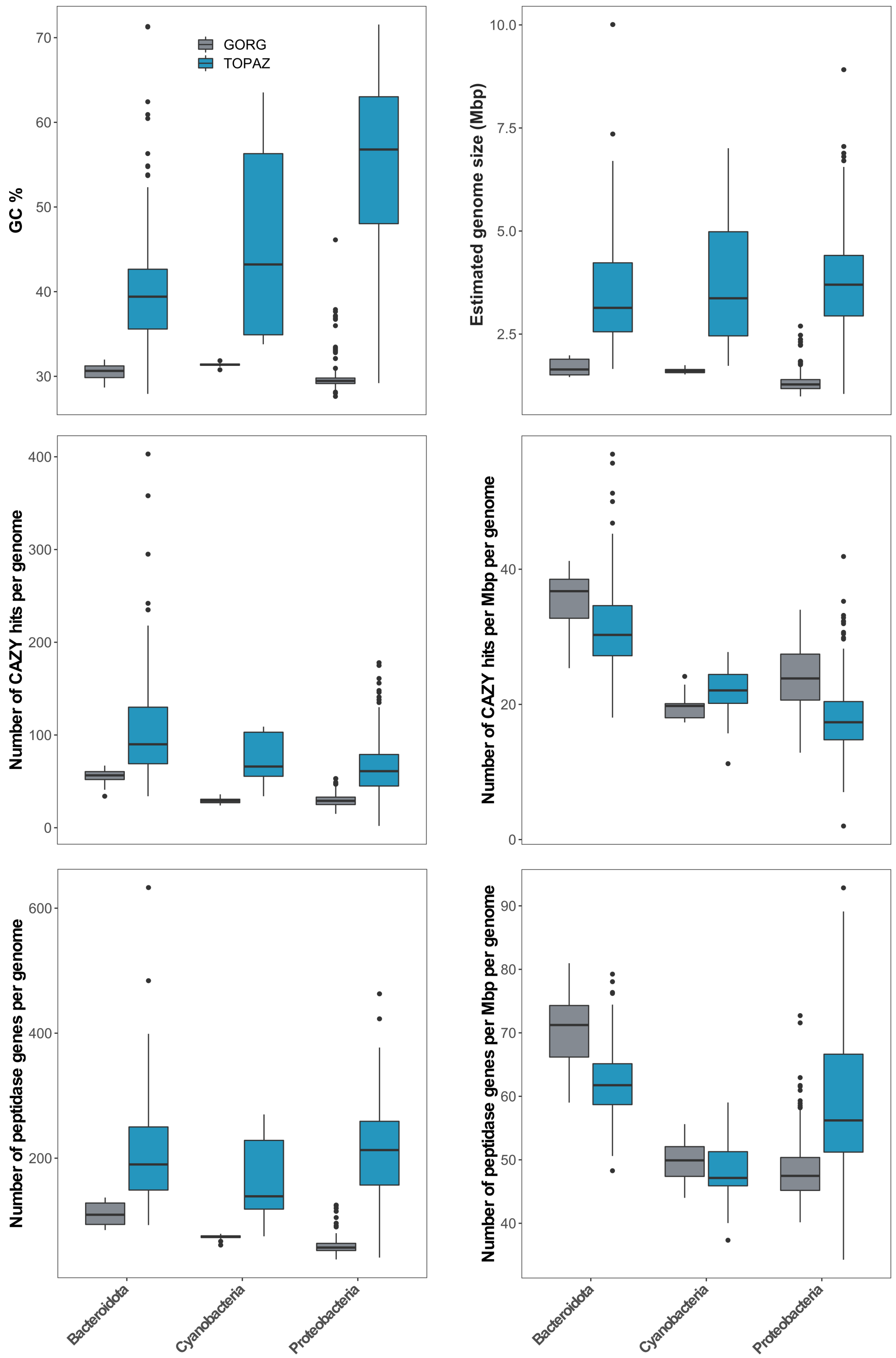


**Figure S22. Comparison of the genomic characteristics between the TOPAZ MAGs and GORG SAGs belonging to the phyla, Bacteroidota, Cyanobacteria and Proteobacteria.** The distributions of GC % content, estimated genome size (Mbp), number of CAZY hits per genome, number of CAZY hits per Mbp per genome, number of peptidase genes per genome and number of peptidase genes per Mbp per genome are presented as box plots.

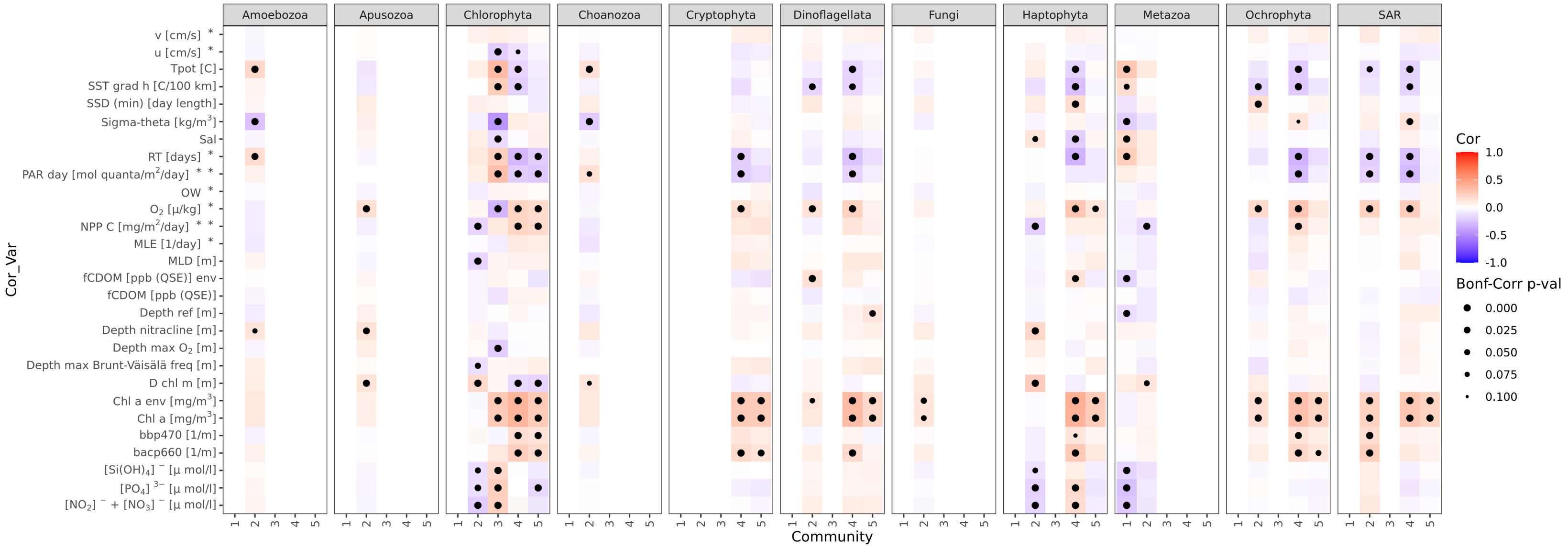


**Figure S23. Extended environmental correlations figure displaying the strength and directions of environmental correlations as tracked by both taxonomic group and network-based community.**

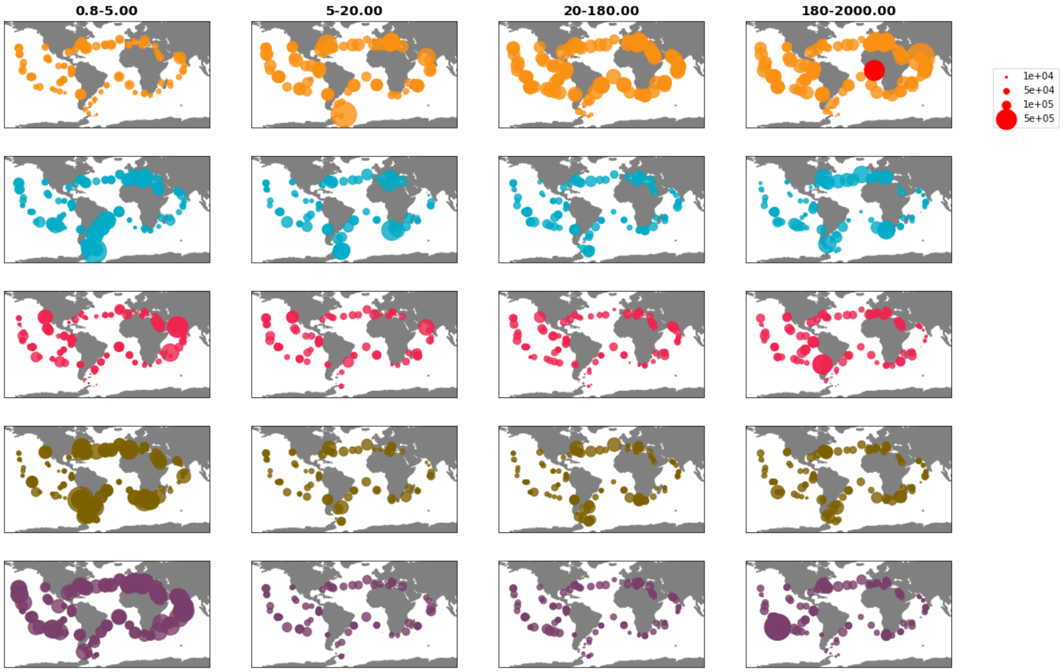


**Figure S24. Distribution of identified communities across *Tara* Oceans Samples. The summed metagenomic abundance (CPM) of each of the seven identified communities from the network analysis (Figure 6) is shown for each size fraction.** The size of the bubble indicates the relative abundance of the community in a given sample.

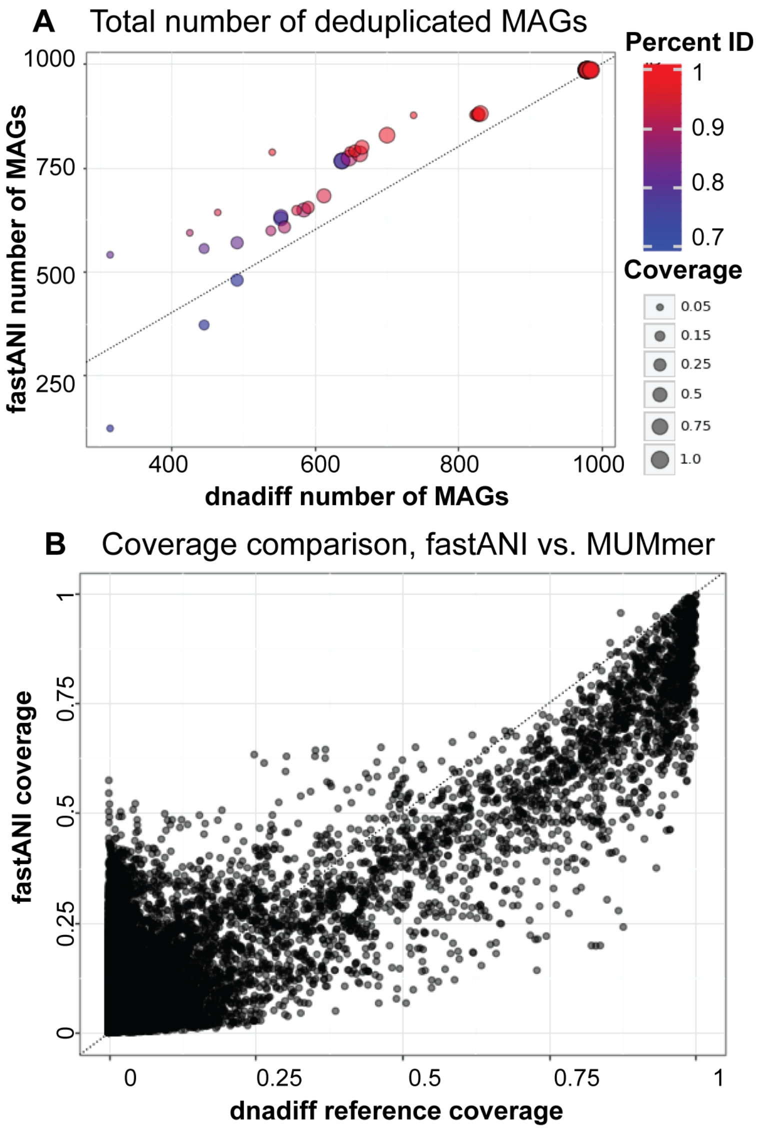


**Figure S25. Comparison of fastANI and dnadiff (MUMmer) approaches for deduplicating MAGs.** A: total number of deduplicated eukaryotic MAGs from fastANI vs. MUMmer; size of points displays the coverage threshold used, while color of points displays the percentage identity threshold used. B: coverage comparison of individual MAGs using fastANI vs. MUMmer.

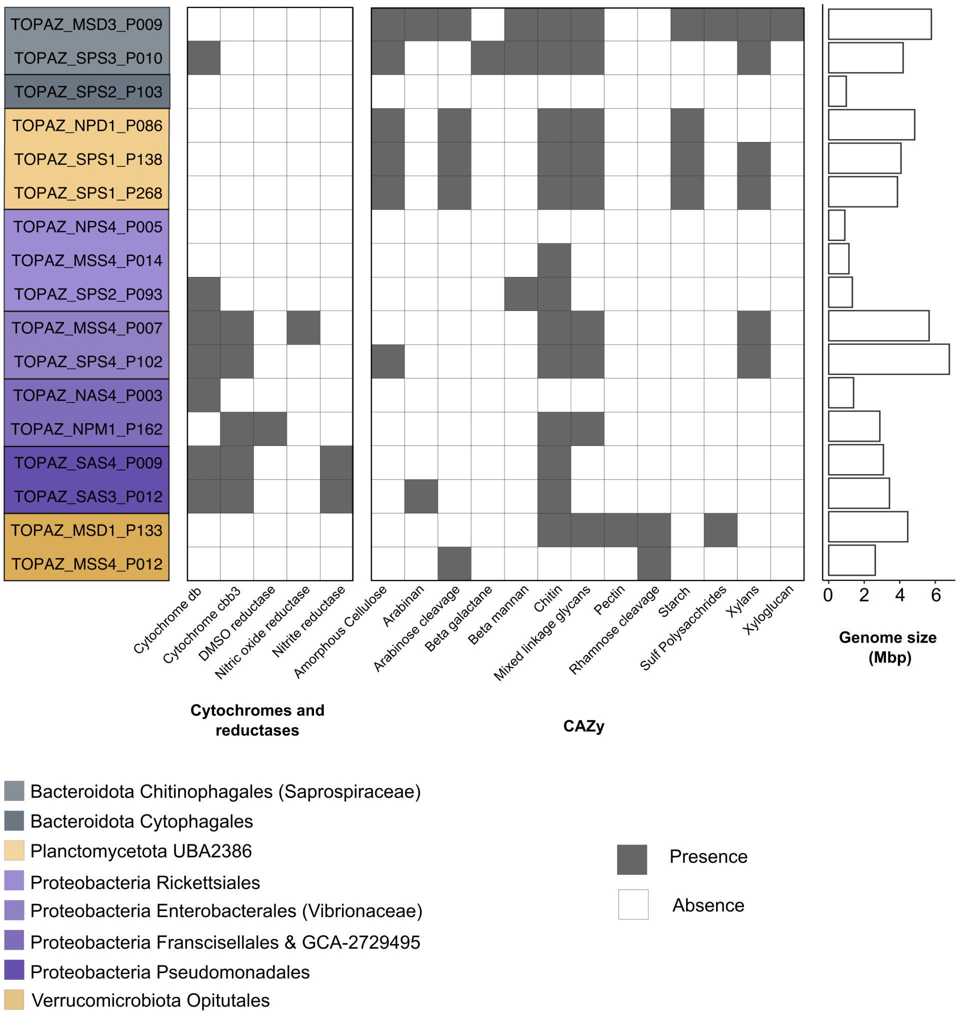


**Figure S26. Genomic characteristics of the Community 1 MAGs.** MAG names are color-coded based on the taxonomic affiliations estimated by GTDB-tk. The presence of the high oxygen affinity cytochromes db ubiquinol oxidase and cbb3, as well as the reductases (involved in anaerobic processes) DMSO family type II, nitric oxide and nitrite are noted with grey in the left panel of the heat map. The presence of genes involved in CAZy pathways for the hydrolysis/degradation of amorphous cellulose, arabinan, arabinose cleavage, beta-galactan, beta-mannan, chitin, mixed-linkage glucans, pectin, rhamnose cleavage, starch, sulf-polysachharides, xylans, xyloglucan are noted with grey in the right panel of the heat map. The estimated size of the genomes is shown as a bar plot.

### References


1. M. R. Olm, C. T. Brown, B. Brooks, J. F. Banfield, *ISME Journal* **11**, 2864 (2017).
2. J. T. Evans, V. J. Denef, *mSphere* **5**, e00971 (2020).
3. G. Marçais, *et al.*, *PLoS Computational Biology* **14**, e1005944 (2018).
4. C. M. Singleton, *et al.*, *Nature Communications* **12**, 1 (2021).
5. E. Bell, *et al.*, *The ISME Journal* pp. 1–11 (2022).
6. P. J. Keeling, *et al.*, *PLOS Biology* **12**, 1 (2014).
7. D. J. Richter, C. Berney, J. F. H. Strassert, F. Burki, d. C. Vargas, *bioRxiv* p. 2020.06.30.180687 (2020).
8. A. Vabalas, E. Gowen, E. Poliakoff, A. J. Casson, *PloS ONE* **14**, e0224365 (2019).
9. L. V. Utkin, *Expert Systems with Applications* **141**, 112978 (2020).
10. L. Collins, G. McCarthy, A. Mellor, G. Newell, L. Smith, *Remote Sensing of Environment* **245**, 111839 (2020).
11. M. Khalilia, S. Chakraborty, M. Popescu, *BMC medical informatics and decision making* **11**, 1 (2011).
12. R. I. Jones, *Freshwater Biology* **45**, 219 (2000).
13. M. Kanehisa, *Protein Science* **28**, 1947 (2019).
14. J. A. Burns, A. A. Pittis, E. Kim, *Nature Ecology & Evolution* **2**, 697 (2018).
15. A. I. Krinos, S. K. Hu, N. R. Cohen, H. Alexander, *Journal of Open Source Software* (2021).
16. T. O. Delmont, *et al.* (2020).
17. C. T. Brown, L. Irber, *JOSS* **1**, 27 (2016).
18. C. Tara Oceans Consortium, P. Tara Oceans Expedition, *Environmental context of all samples from the Tara Oceans Expedition (2009-2013), about water column features* (PANGAEA, 2016). In: Tara Oceans Consortium, C; Tara Oceans Expedition, P (2016): Registry of all samples from the Tara Oceans Expedition (2009-2013). PANGAEA, https://doi.org/10.1594/PANGAEA.859953.
19. F. d'Ovidio, S. D. Monte, S. Alvain, Y. Dandonneau, M. Levy, *Proceedings of the National Academy of Sciences* **107**, 18366 (2010).
